# The active repertoire of *Escherichia coli* peptidoglycan amidases varies with physiochemical environment

**DOI:** 10.1101/2020.10.19.344754

**Authors:** Elizabeth A. Mueller, Abbygail G. Iken, Mehmet Ali Öztürk, Mirko Schmitz, Barbara Di Ventura, Petra Anne Levin

**Affiliations:** Department of Biology, Washington University in St. Louis, St. Louis, Missouri, USA; Signalling Research Centers BIOSS and CIBSS, University of Freiburg, 79104, Freiburg, Germany; Institute of Biology II, Faculty of Biology, University of Freiburg, 79104, Freiburg, Germany

**Keywords:** Amidases, peptidoglycan, pH, cytokinesis, morphogenesis

## Abstract

Nearly all bacteria are encased in a peptidoglycan cell wall, an essential crosslinked matrix of polysaccharide strands and short peptide stems. In the Gram-negative model organism *Escherichia coli*, more than forty cell wall synthases and autolysins coordinate the growth and division of the peptidoglycan sacculus in the periplasm. The precise contribution of many of these enzymes to cell wall metabolism remains unclear due to significant apparent redundancy, particularly among the cell wall autolysins. *E. coli* produces three major LytC-type-*N*-acetylmuramoyl-*L*-alanine amidases, which share a role in separating the newly formed daughter cells during cytokinesis. Here, we reveal two of the three amidases exhibit growth medium-dependent changes in activity. Specifically, we report acidic growth conditions stimulate AmiB—and to a lesser extent, AmiC—activity. Combining computational and genetic analysis, we demonstrate that low pH-dependent stimulation of AmiB requires three periplasmic amidase activators: EnvC, NlpD, and YgeR. Altogether, our findings support overlapping, but not redundant, roles for the *E. coli* amidases in cell separation and illuminate the physiochemical environment as an important mediator of cell wall enzyme activity.

**IMPORTANCE:** Penicillin and related β-lactam antibiotics targeting the bacterial cell wall synthesis are among the most commonly prescribed antimicrobials worldwide. However, rising rates of antibiotic resistance and tolerance jeopardize their continued clinical use. Development of new cell wall active therapeutics, including those targeting cell wall autolysins, has been stymied in part due to high levels of apparent enzymatic redundancy. In this study, we report a subset of *E. coli* amidases involved in cell separation during cell division are not redundant and instead are preferentially active during growth in distinct pH environments. Specifically, we discover *E. coli* amidases AmiB and AmiC are activated by acidic pH. Three semi-redundant periplasmic regulators—NlpD, EnvC, and YgeR—collectively mediate low pH-dependent stimulation of amidase activity. This discovery contributes to our understanding of how the cell wall remains robust across diverse environmental conditions and reveals opportunities for the development of condition-specific antimicrobial agents.

## INTRODUCTION

Nearly all bacteria are surrounded by peptidoglycan (PG) cell wall, which protects cells from osmotic rupture and confers cell shape. PG consists of glycan strands of repeating disaccharides of *N*-acetylglucosamine and *N*-acetylmuramic acid sugars. Adjacent glycan strands are crosslinked to one another through short peptide stems to form a continuous matrix, referred to as the sacculus. The growth and division of the sacculus must be coordinated with cell replication to prevent lethal breaches in the PG. Accordingly, small molecules that disrupt this process, including β-lactam and glycopeptide antibiotics, are among the oldest and most effective weapons against bacterial pathogens.

The reactions involved in PG synthesis span both the cytoplasm and periplasm in Gram-negative bacteria. Cytoplasmic enzymes generate lipid-linked disaccharide precursors, which are translocated to the outer leaflet of the plasma membrane by the flippase MurJ (1). In the periplasm, two classes of enzymes assemble and modify the new PG material: PG synthases and autolysins. PG synthases polymerize and crosslink glycan strands. PG autolysins, on the other hand, exhibit diverse biochemical activities, and their cumulative activity can break nearly every bond in the PG sacculus (2). The activities of PG synthases and autolysins must be tightly coupled to one another and to the cell cycle to preserve PG integrity (3). Spatiotemporal regulation of these enzymes is particularly important during cytokinesis, which requires concomitant synthesis and hydrolysis of cell wall material in order to form and separate the new daughter cells.

The precise cellular function of many periplasmic cell wall enzymes remains unclear due to significant enzymatic redundancy, particularly among PG autolysins (4–8). As many as eight enzymes are capable of catalyzing each autolytic reaction in *E. coli* cell wall metabolism (9). In some cases, apparently redundant PG enzymes can be distinguished by differences in substrate specificity (10, 11), subcellular localization (12–14), and interaction partners (13, 15–18). Additionally, recent work from our group and others revealed the activity of certain “redundant” PG synthases and autolysins changes based on the physiochemical properties of the growth medium (19–25). We discovered two semi-redundant *E. coli* PG synthases, PBP1a and PBP1b, are preferentially required for PG integrity in opposing pH environments, in part due to pH-dependent differences in enzymatic activity (21). Collectively, this work suggests the active repertoire of periplasmic PG enzymes is modular with respect to the growth environment (9, 26), with some enzymes only active in a subset of growth conditions (19–22, 24). We hypothesize these enzymes, henceforth referred to as condition-dependent “specialists”, ensure robustness in the periplasmic steps of PG metabolism, which are exposed to the physiochemical properties of the extracellular medium (27–29). In support of this model, pH specialist enzymes have been identified in every major class of *E. coli* PG autolysins except one—the amidases (19–24).

Amidases cleave the peptide stem from the glycan backbone of PG, producing stem-less or “denuded” glycans (30) (**Fig. 1A**). This reaction is required to separate newly formed daughter cells following cross-wall synthesis during cytokinesis. *E. coli* produces three major LytC-type-*N*-acetylmuramoyl-*L*-alanine amidases with overlapping roles in cell separation: AmiA, AmiB, and AmiC (**Fig. 1B**). Several pieces of evidence support functional redundancy between these three enzymes. Cells separate normally in the absence of any individual amidase, but loss of multiple amidases leads to the formation of chains of unseparated cells that retain distinct cytoplasmic compartments but share an outer membrane (5, 31). At the same time, cytological and interaction studies indicate these enzymes have overlapping, but not necessarily redundant, activities. Notably, the amidases differ in subcellular localization profiles. AmiB and AmiC are enriched at the invaginating septum during cross-wall synthesis, while AmiA is uniformly distributed around the cell periphery throughout the cell cycle (14, 32). Moreover, unique interaction partners stimulate amidase activity. While all three amidases possess an autoinhibitory helix that prevents aberrant autoactivation (33), different LtyM family protein activators are required to relieve autoinhibition and license their cognate amidases’ cleavage activity at midcell during cytokinesis (34). The periplasmic protein EnvC activates AmiA and AmiB, whereas the outer membrane-anchored NlpD promotes AmiC activity (35–38). Whether the amidases or their regulators also exhibit condition-dependent differences in activity remains unknown.

**Fig. 1.**
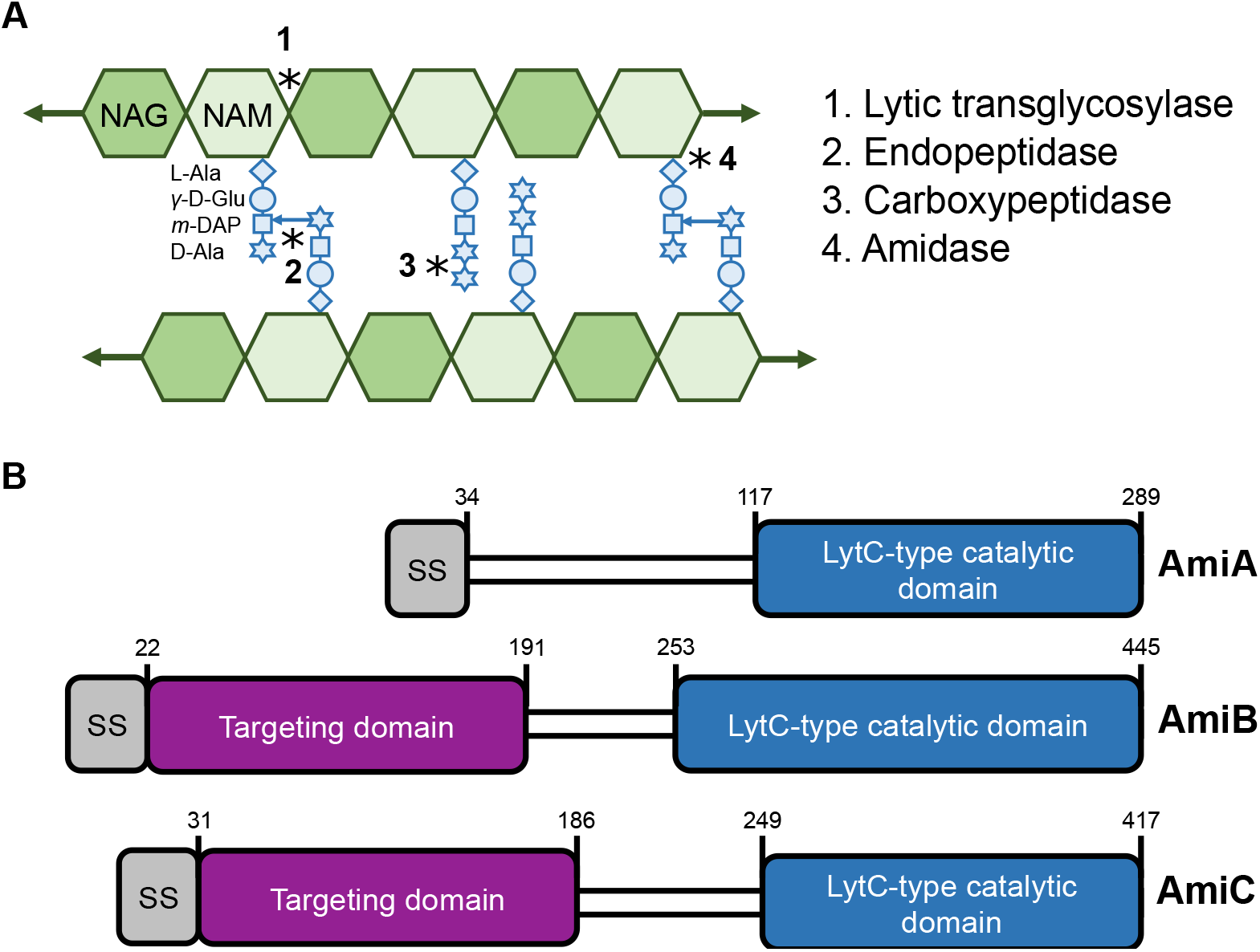
*E. coli* produces three LytC-type-*N*-acetylmuramoyl-*L*-alanine amidases. A) A model of the peptidoglycan sacculus. *N*-acetylglucosamine (NAG) and *N*-acetylmuramic acid (NAM) sugars conjoined by a β-(1,4) linkage form glycan strands. Peptide stems emanate from the NAM sugar and may be crosslinked to adjacent glycan strands. Peptidoglycan autolysins cleave the bonds indicated by asterisks. B) *E. coli* produces three LytC-type-*N*-acetylmuramoyl-*L*-alanine amidases involved in cell separation. Schematic depicts signal sequence (grey), septum targeting domain (purple), and LytC-type catalytic domain (blue). Note that AmiA and AmiC are targeted through the periplasm through the twin arginine translocation (Tat) protein export pathway, while AmiB is targeted through the Sec protein translocation pathway.

Here, we evaluated the impact of environmental pH on *E. coli* amidase activity. By comparing the morphology of variants producing a single amidase at pH 6.9 and 5.2, we report that AmiB and AmiC have heightened activity in acidic medium, whereas AmiA activity is insensitive across this pH range. AmiB, in particular, is unable to support cell separation in neutral medium but becomes sufficient for cell separation in acidic conditions. Genetic and computational analysis indicate AmiB stimulation in acidic medium is due to promiscuous activation by EnvC, NlpD, and YgeR, a third LtyM family protein specifically involved in cell separation in acidic conditions. Altogether, our work supports a central role for the extracellular environment in dictating the active repertoire of cell wall enzymes and regulatory proteins.

## RESULTS

### Acidic pH alleviates cell separation defect of amidase mutants

To assess the impact of pH on the activity of individual amidases, we compared the growth and morphological phenotypes of cells deficient for one or a combination of the three major *E. coli* LytC-type-*N*-acetylmuramoyl-*L*-alanine amidases—AmiA, AmiB, and AmiC. Deletion alleles were sourced from the Keio deletion collection (39) with the exception of Δ*amiB* (a gift from Daniel Kahne, (40)). We were unable to verify the Keio version of Δ*amiB* by Sanger sequencing and hence elected not to use it. All alleles were transduced into the wild-type parental strain MG1655 (**SI Appendix, Table S1A**). We cultured the resulting bacterial strains to steady state in LB medium buffered to pH 5.2 or 6.9 and then imaged the cells by phase contrast microscopy (see **Materials and Methods**). We recorded the growth rate and chain length for each mutant as a function of environmental pH (**SI Appendix, Table S2**).

While none of the single amidase mutants exhibited pH-dependent changes in growth rate or morphology (**SI Appendix, Fig. S1 and Table S2**), analysis of the double mutants indicates that two of *E. coli’s* three amidases, AmiB and AmiC, are sensitive to pH. Double mutants encoding only *amiB* or *amiC* (*amiB*^+^ Δ*amiAC* or *amiC*^+^ Δ*amiAB*, respectively) displayed dramatic pH-dependent chaining phenotypes (**Fig. 2; SI Appendix, Table S2**). Consistent with previous studies (5, 40), each of these strains formed chains at pH 6.9, albeit to differing extents. *amiB*^+^ Δ*amiAC* cells formed long chains of 25 ± 1 cells in length, while *amiC*^+^ Δ*amiAB* formed short chains of 2 to 4 cells in length. As expected, normal cell separation could be restored by ectopic expression of the missing amidase alleles from a plasmid (**SI Appendix, Fig. S3**). More surprisingly, culturing the double mutants in acidic medium was also sufficient to reduce chaining by up to 86% (*amiB*^+^ Δ*amiAC*, mean of 4 ± 1 cells per chain; mean of *amiC*^+^ Δ*amiAB*, 1 ± 1 cell per chain). The dramatic reduction in chaining was accompanied by only a minimal reduction in growth rate (**SI Appendix, Table S2**).

**Fig. 2.**
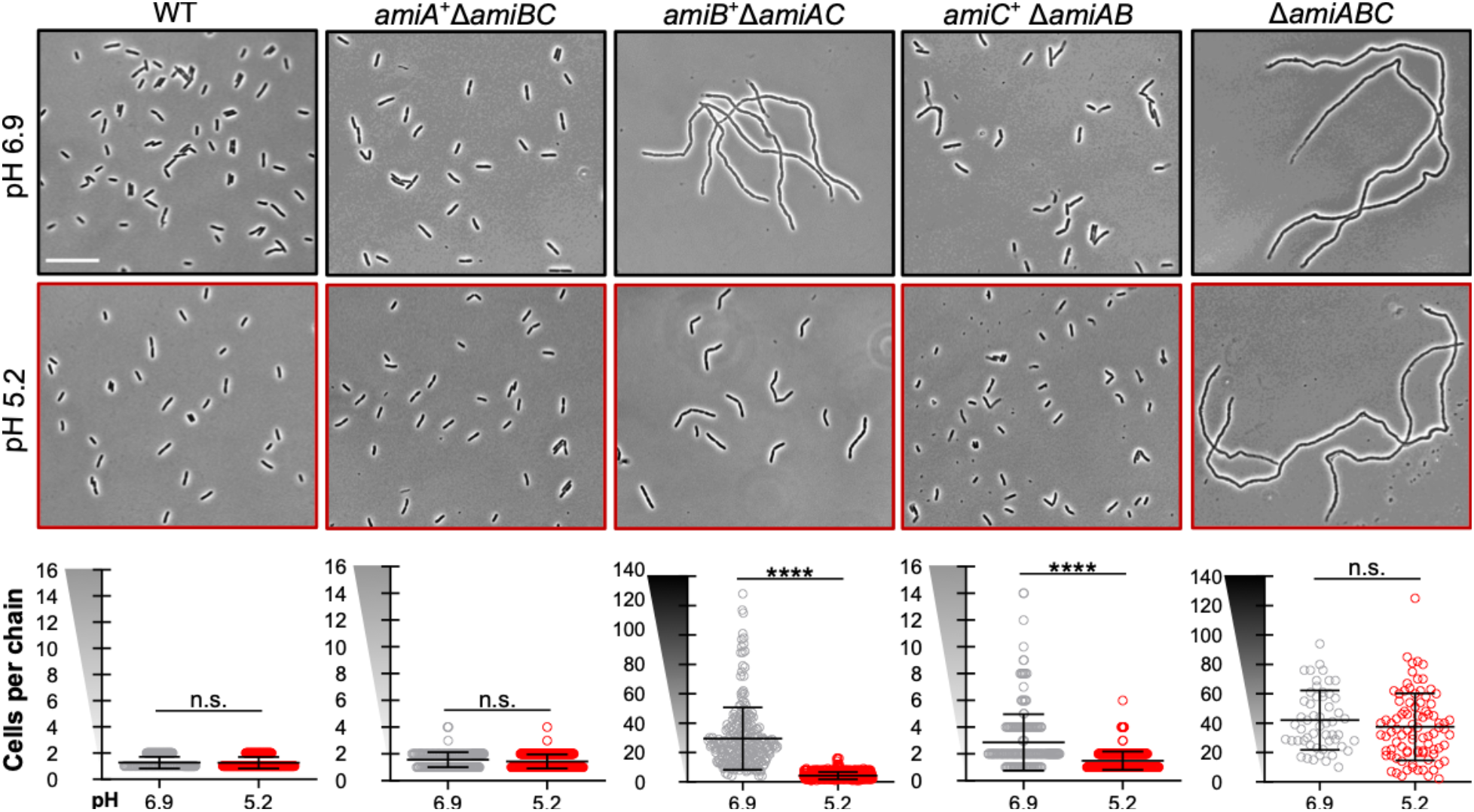
Acidic pH alleviates cell separation defect of amidase mutants. Representative micrographs of double [Δ*amiAB* (EAM1379), Δ*amiAC* (EAM927), and Δ*amiBC* (EAM1381)] and triple amidase (Δ*amiABC*, EAM1385) mutants during steady state growth in LB medium buffered to pH 6.9 or 5.2 compared to the parental strain (MG1655). Cells were grown to mid-exponential phase (OD_600_ ~ 0.2-0.6) at the indicated pH, sub-cultured into the same medium at an OD_600_ ~ 0.005, then sampled and fixed at OD_600_ ~ 0.1-0.2 for microscopy. Scale bar denotes 20 μm. Quantification for each mutant is shown below the corresponding set of micrographs. Points indicate individual chain length measurements from at least two independent experiments, and error bars denote standard deviation. Y-axis gradient emphasizes the difference in scale used between mutants. Significance was determined by Kolmogorov-Smirnov test with a threshold of statistical significance set to p < 0.01.

To validate the findings for the *amiB*^+^ Δ*amiAC* mutant, we took advantage of mutants defective in the twin-arginine translocation (Tat) protein export pathway (Δ*tatB* and Δ*tatC*), which is required for the secretion of AmiA and AmiC into the periplasm (14, 41). Consistent with previous reports (14, 41), Δ*tatB* and Δ*tatC* mutants were phenotypically identical to the *amiB*^+^ Δ*amiAC* mutant at pH 6.9. Likewise, growth in acidic medium dramatically reduced the chain length of Δ*tatB* and Δ*tatC* mutants (**SI Appendix, Fig. S2**), highlighting acid-dependent stimulation of AmiB activity. These findings reinforce a model in which AmiB is largely sufficient for cell separation during growth in acidic, but not neutral, pH conditions.

In contrast to AmiB and AmiC, AmiA appears sufficient for normal cell separation during growth in both neutral and acidic conditions. An *amiA*^+^ Δ*amiBC* double mutant separated normally when cultured at pH 6.9 or 5.2. Conversely, the Δ*amiABC* triple mutant formed long chains of unseparated cells at both acidic and neutral pH (pH 6.9, mean of 44 ± 2 cells per chain; pH 5.2, mean of 37 ± 5 cells per chain) (**SI Appendix, Table S2**).

### Acidic pH restores cell envelope integrity of amidase mutants

Defects in cell separation are typically associated with increases in cell envelope permeability, rendering cells sensitive to hydrophobic compounds, detergents, and certain antibiotics (38, 41–43). We reasoned that since pH influences cell separation, the amidase mutants may also exhibit differential envelope permeability across pH conditions. To assess cell envelope permeability, we compared the amidase double mutants’ sensitivity to the commonly used detergent Triton-X during growth in neutral (pH 6.9) or acidic (pH 5.2) medium. Relative sensitivity was assessed by determining the minimum inhibitory concentration of Triton-X for each strain.

Supporting the pH sensitivity of AmiB and AmiC, *amiB*^+^ Δ*amiAC* and *amiC*^+^ Δ*amiAB* strains were up to 64-fold more resistant to Triton-X when cultured in acidic medium compared to neutral medium (**Fig. 3**). In contrast, wild-type cells and *amiA*^+^ Δ*amiBC* mutants, both of which exist primarily as individual cells (**Fig. 2**), were highly resistant to Triton-X at both pH conditions (MIC of 10% Triton-X). The Δ*amiABC* triple mutant remained highly sensitive to the detergent at both pH conditions (MIC of 0.039% Triton-X). Altogether, our data indicate that acidic pH partially alleviates the cell separation defect and cell envelope permeability of Δ*amiAC* and Δ*amiAB* double mutants, presumably through activation of AmiB or AimC.

**Fig. 3.**
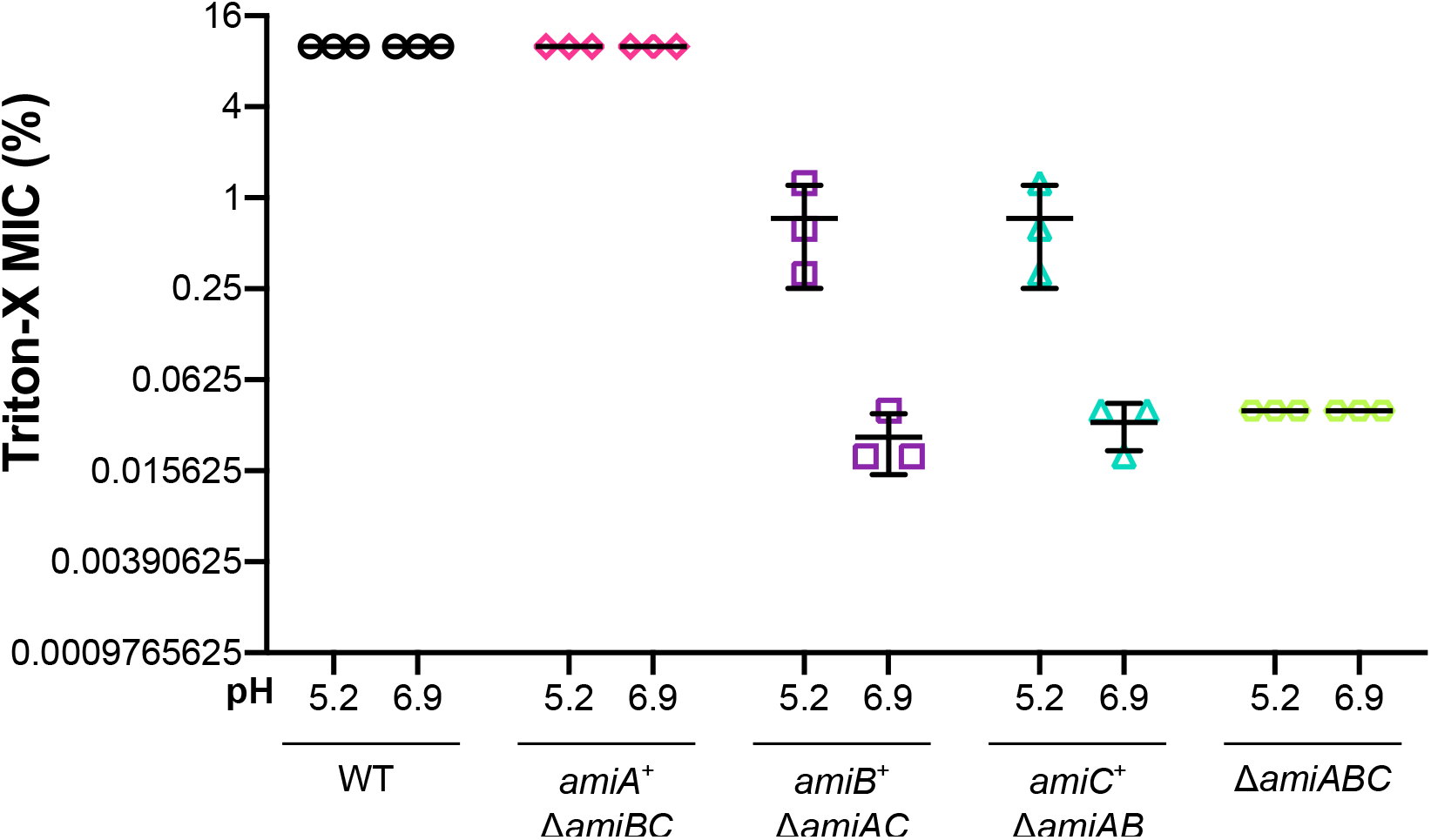
Acidic pH restores cell envelope integrity of amidase mutants. Comparison of the minimum inhibitory concentration (MIC) of Triton-X for indicated mutants when cultured in buffered LB medium at pH 6.9 or 5.2 [wild type (MG1655), Δ*amiAB* (EAM1379), Δ*amiAC* (EAM927), Δ*amiBC* (EAM1381), and Δ*amiABC* (EAM1385)]. Cells were grown to mid-exponential phase (OD_600_ ~ 0.2-0.6) in LB medium at pH 6.9, then sub-cultured into 96-well plates to 1×10^5^ CFU/mL in LB medium buffered to either pH 6.9 or 5.2 with varying percentages of Triton-X detergent. Cells were grown with aeration at 37 °C for 20 hours before MIC was determined by visual inspection. Error bars denote standard deviation.

### Acidic pH stimulates AmiB and AmiC activity *in vivo*

Our genetic data are consistent with two models. First, AmiB and AmiC activity is sufficiently enhanced at low pH to support cell separation the absence of other amidases. Alternatively, cell separation in acidic conditions may be mediated in part by an unknown amidase or by another class of cell wall autolysins. Endopeptidases and lytic transglycosylases also play minor roles in cell separation under standard culture conditions (6, 43, 44). Consistent with the former model, cells deficient for all three amidases (Δ*amiABC*) produced long chains of similar cell lengths and displayed equivalent sensitivity to Triton-X when cultured under either pH condition (**Fig. 2, 3; SI Appendix, Table S2**). Moreover, overexpression of *amiB* from an arabinose-inducible promoter restored cell separation to *amiB*^+^ Δ*amiAC* mutants at pH 6.9, suggesting that increases in basal AmiB activity are sufficient to alleviate chaining (**SI Appendix, Fig. S3**).

We employed two additional independent approaches to examine the influence of pH on AmiB and AmiC activity more directly: 1) assessing the impact of overexpression of individual amidase genes in wild-type cells, and 2) comparing the production of denuded glycan through cytological analysis at neutral and acidic pH. The first approach takes advantage of the observation that amidase activity is frequently associated with cell lysis and morphological defects (33, 45, 46). If acidic growth medium enhances AmiB and AmiC activity, lowering the pH should exacerbate any lytic or morphological phenotypes associated with their overproduction. We therefore overexpressed *amiA, amiB*, and *amiC* from an arabinose-inducible promoter on a plasmid (pBAD) in otherwise wild-type cells and compared cell growth and morphology across pH conditions.

Supporting enhanced AmiB activity, acidic pH exacerbated the toxicity of *amiB* overexpression. At high concentrations of arabinose (0.4%), AmiB overproduction stimulated cell lysis and rounding under both pH conditions (**SI Appendix, Fig. S4A, B**), phenotypes indicative of disrupted cell wall homeostasis. Consistent with acid-mediated activation, AmiB production was lytic at pH 5.2, but not pH 6.9, at ten-fold lower concentrations of inducer (0.04%) (**Fig. 4**). Unlike *amiB*, cells overexpressing *amiA* or *amiC* were indistinguishable from those harboring the empty vector control at either pH value (**SI Appendix, Fig. S5C-E**).

**Fig. 4.**
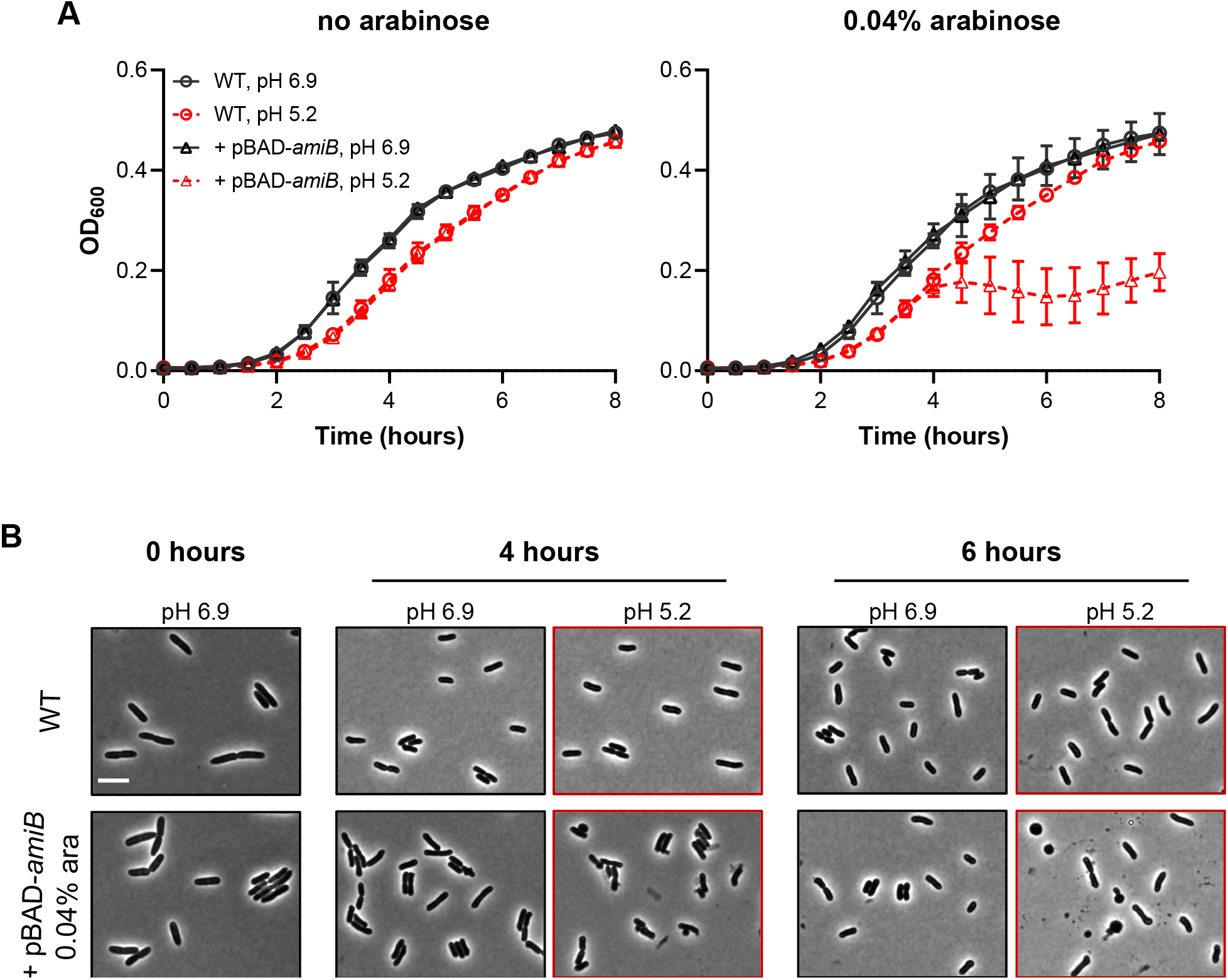
AmiB overproduction induces lysis and cell rounding in acidic medium. A) Growth curves of wild-type (MG1655) cells overexpressing *amiB* from a plasmid in the presence or absence of inducer (0.04% arabinose) when cultured at pH 6.9 or 5.2. Cells were grown to mid-exponential phase (OD_600_ ~ 0.2-0.6) in LB medium at pH 6.9, then sub-cultured into 96-well plates at an OD_600_ ~ 0.005 in LB medium buffered to either pH 6.9 or 5.2. Cells were grown with aeration at 37 °C, and the optical density was measured every 10 minutes. Curves represent the average ± standard deviation of three independent experiments. B) Representative micrographs of cell morphology after two, four, or six hours of growth with inducer. Scale bar denotes 5 μm.

To further assess the impact of pH on amidase activity and exclude indirect effects by another class of PG autolysins, we leveraged the finding that the products of amidase activity—”denuded” glycans lacking stem peptides—are specifically recognized by “sporulation-related repeat” (SPOR) domain-containing proteins. These proteins include the *E. coli* cell division proteins DamX, DedD, and FtsN, which rely on the SPOR domain to localize to midcell during cytokinesis (47, 48). Since amidase activity is concentrated at sites of septation, there should be an increase in denuded glycans at medial and polar positions on the surface of growing cells. Reasoning that midcell binding of a fluorescent protein-SPOR domain fusion should reflect levels of septal amidase activity *in vivo*, we incubated purified His_6_-GFP-DamX^SPOR^—which binds denuded glycans in isolated sacculi (47) —with fixed wild-type, *amiA*^+^ Δ*amiBC*, *amiB*^+^ Δ*amiAC*, or *amiC*^+^ Δ*amiAB* cells (see **Materials and Methods; SI Appendix, Fig. S5A**). Notably, while cells were cultured at either neutral or acidic pH prior to fixation, the SPOR domain binding assay was performed on fixed cells equilibrated to pH 7.4 to ensure uniform reactions conditions.

Consistent with SPOR domain binding serving as a *bone fide* reporter of denuded glycans in fixed cells, His_6_-GFP-DamX^SPOR^ fluorescence was concentrated at midcell in wild-type *E. coli* cultured at neutral and acidic pH (**Fig. 5A; SI Appendix, Figure S6B,C**). As expected, we did not observe midcell enrichment of His_6_-GFP-DamX^SPOR^ localization in the Δ*amiABC* mutant (47, 49) (**SI Appendix, Fig. S5B,C**). We did, however, observe nonspecific His_6_-GFP-DamX^SPOR^ binding at all positions along the cell body of the *ΔamiABC* mutant, likely due to the enhanced cell envelope permeability in this mutant (**Fig. 3**). Conversely, medial fluorescence from His_6_-GFP-DamX^SPOR^ was enhanced four-fold over wild type in a strain that lacks six lytic transglycosylases (Δ6LT) (**SI Appendix, Figure S6B,C**); this strain was previously shown to have elevated levels of denuded glycans (47), possibly due to reduced glycan processing (49). Importantly, chain length is not necessarily correlated with His_6_-GFP-DamX^SPOR^ intensity at the septum, as the Δ6LT mutant forms chains 2-8 cells in length (**SI Appendix, Figure S6D**).

**Fig. 5.**
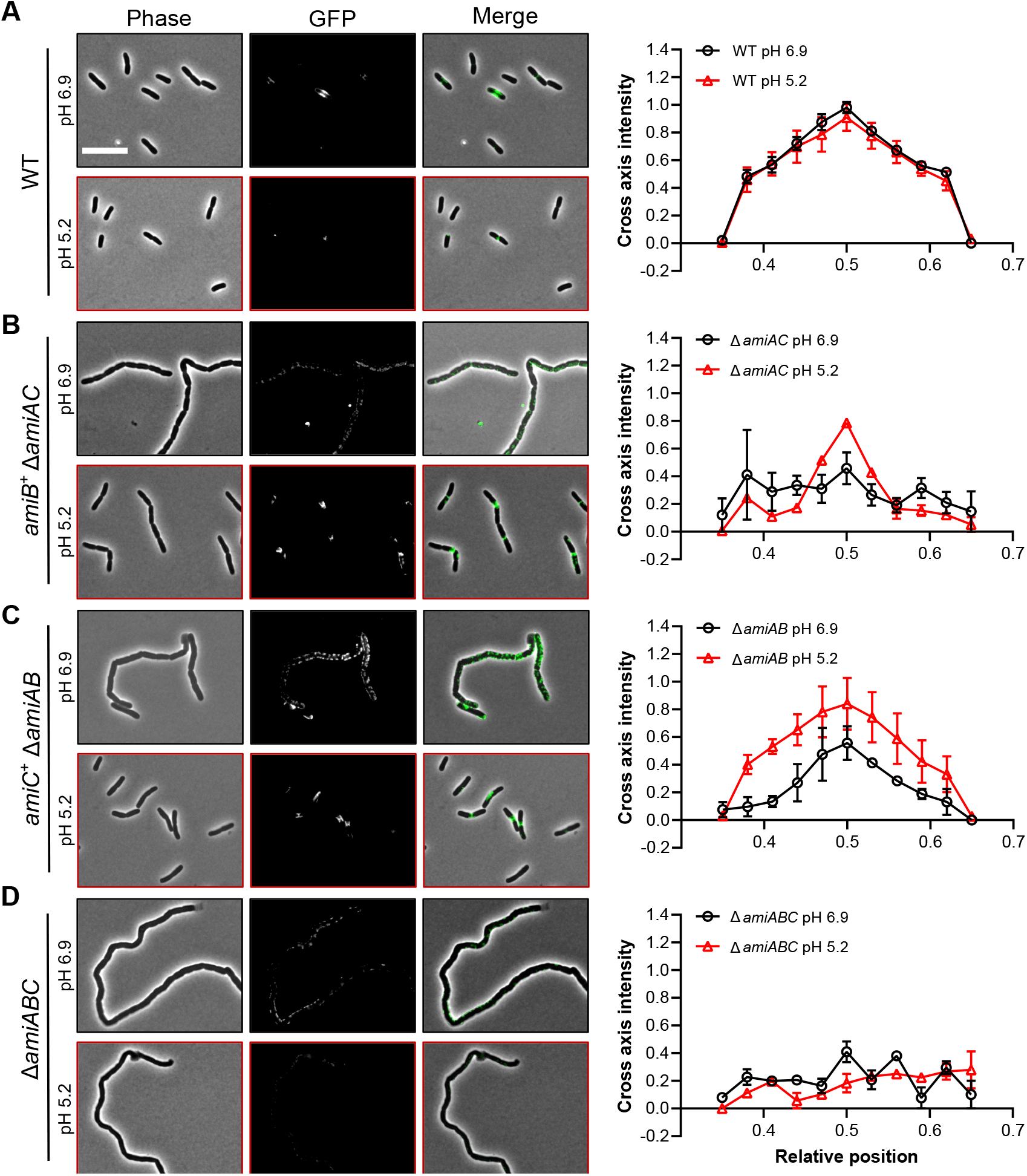
SPOR domain binding assay indicates elevated AmiB and AmiC activity at midcell in acidic medium. Representative micrographs and normalized intensity profiles for wild type (MG1655, A), Δ*amiAC* (EAM927, B), Δ*amiBC* (EAM1381, C), and Δ*amiABC* (EAM1385, D) after labeling with His_6_-GFP-DamX^SPOR^ protein. Cells were cultured to steady state in LB medium (pH 6.9 or 5.2) and then fixed at OD_600_ ~ 0.1-0.2. Fixed cells were immediately labeled with 100 ng/mL of purified His_6_-GFP-DamX^SPOR^ protein for 30 minutes on ice, washed, and then visualized by fluorescence microscopy. Fluorescence intensity profiles were generated from at least 50 cells across two biological replicates and normalized to the maximum intensity of the wild type at pH 6.9. Error bars represent standard deviation. Scale bar denotes 10 μm.

In support of elevated AmiB and AmiC activity at low pH, *amiB*^+^ Δ*amiAC* or *amiC*^+^ Δ*amiAB* strains cultured at pH 5.2 prior to fixation displayed a ~50% increase in His_6_-GFP-DamX^SPOR^ at midcell relative to their counterparts cultured at 6.9 (**Fig. 5B, C**). This phenotype was particularly stark for the *amiB*^+^ Δ*amiAC* cells, which only exhibited nonspecific probe binding in neutral medium (**Fig. 5B**). Conversely, wild-type cells and cells producing only AmiA (Δ*amiBC*) bound GFP-DamX^SPOR^ with similar affinities at midcell regardless of whether they were cultured at pH 6.9 or 5.2 (**Fig. 5A; SI Appendix, Fig. S5E**). Overall, our overexpression and cytological studies support a model in which low pH stimulates AmiB and AmiC amidase activity to promote cell separation.

### pH-dependent AmiB and AmiC activation is partially independent of EnvC and NlpD

Amidase activity is spatiotemporally controlled *in vivo* through direct interaction with the LytM-domain proteins EnvC and NlpD. The periplasmic protein EnvC (50, 51) and outer membrane lipoprotein NlpD (38) are required for efficient amidase activity *in vitro* (36) and *in vivo* (37) and restrict amidase activation to the midcell after the onset of cytokinesis. At neutral pH, EnvC specifically stimulates AmiA and AmiB, while NlpD is responsible for AmiC activation (36) (**Fig. 6A**), although cross-activation has been observed in some genetic backgrounds (33, 38). In accordance with their requirement for amidase activation, cells defective for EnvC and NlpD production phenotypically mimic an Δ*amiABC* mutant and form long chains of unseparated cells under standard culture conditions (37). To assess whether either one of these regulators mediated AmiB or AmiC activation in acidic medium, we compared cell chaining of an Δ*nlpDenvC* mutant across pH conditions.

**Fig. 6.**
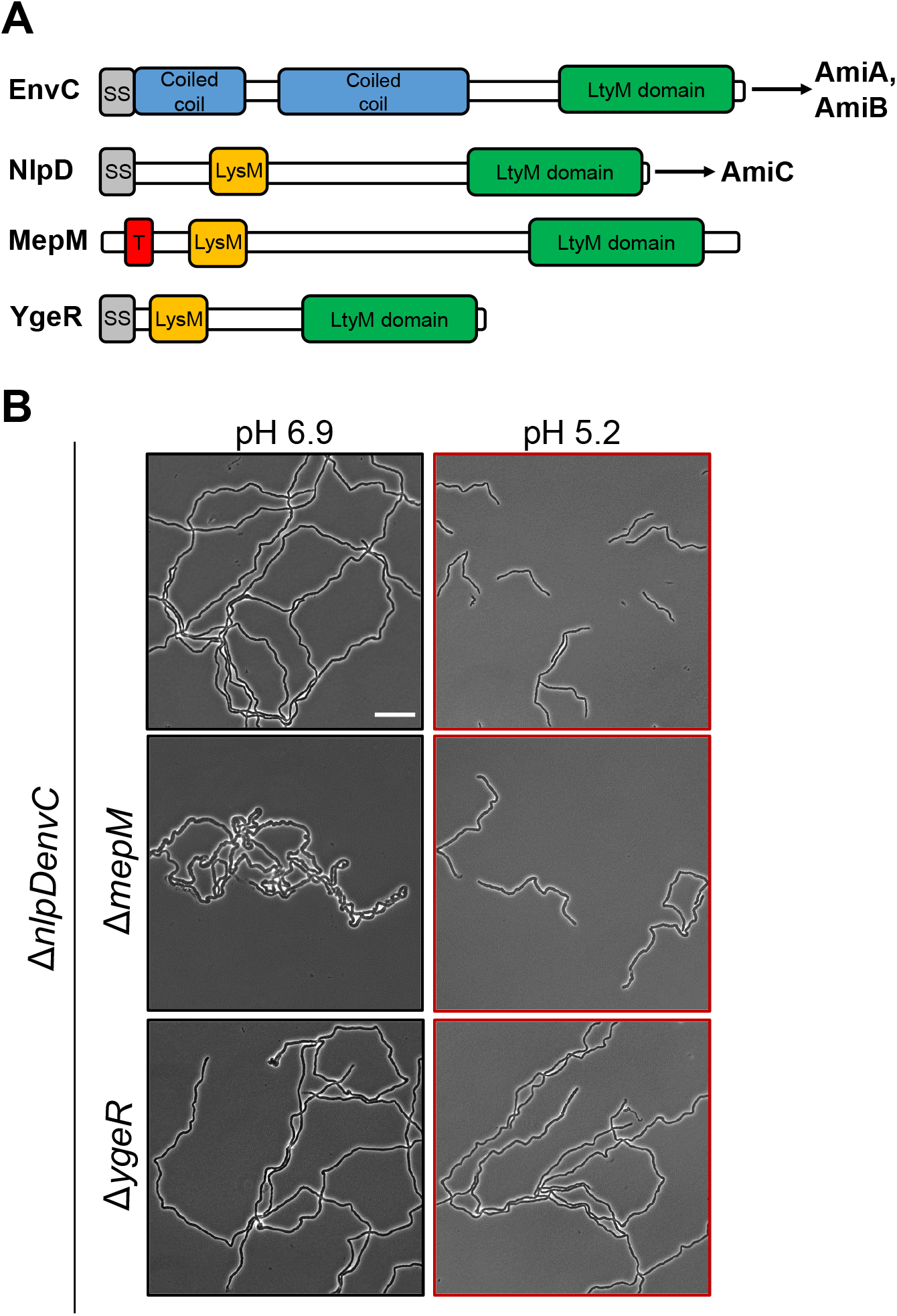
LtyM-domain protein YgeR contributes to cell separation at acidic pH. A) *E. coli* produces four LytM-domain proteins: EnvC, NlpD, MepM, and YgeR. Schematic depicts conserved and unique domains among the proteins, including the signal sequence (grey), coiled coil regions (blue), LysM domain (orange), and the conserved LtyM domain (green). The current model suggests EnvC specifically activates AmiA and AmiB, while NlpD specifically activates AmiC. B) Representative micrographs for Δ*nlpDenvC* (EAM1372), Δ*nlpDenvCyebA* (EAM1477), and Δ*nlpDenvCygeR* (EAM1521) during steady state growth in LB medium buffered to pH 6.9 or 5.2. Cells were grown to mid-exponential phase (OD_600_ ~ 0.2-0.6) at the indicated pH, sub-cultured into the same medium at an OD_600_ ~ 0.005, then sampled at OD_600_ ~ 0.1-0.2 for live cell microscopy. Scale bar denotes 20 μm.

Our data indicate acidic pH stimulates the amidases through a mechanism partially independent of EnvC and NlpD. At neutral pH, the Δ*nlpDenvC* mutant formed chains with an average of 45 ± 3 cells (**Fig. 6B, 7A; SI Appendix, Table S2**), similar to the chain length observed for the Δ*amiABC* triple mutant (**Fig. 2**). However, in contrast to the pH-insensitive Δ*amiABC* triple mutant, the average chain length for the Δ*nlpDenvC* mutant was reduced by 75% to 11 ± 1 cells per chain during growth in acidic medium (**Fig. 6B, 7A; SI Appendix, Table S2)**. The reduction in chaining in acidic medium was dependent on *amiB* and *amiC*, but not *amiA* (**SI Appendix, Fig. S6**). Altogether, the retention of pH sensitivity by the Δ*nlpDenvC* mutant suggests that AmiB and AmiC activation is partially independent of EnvC or NlpD during growth at low pH.

**Fig. 7.**
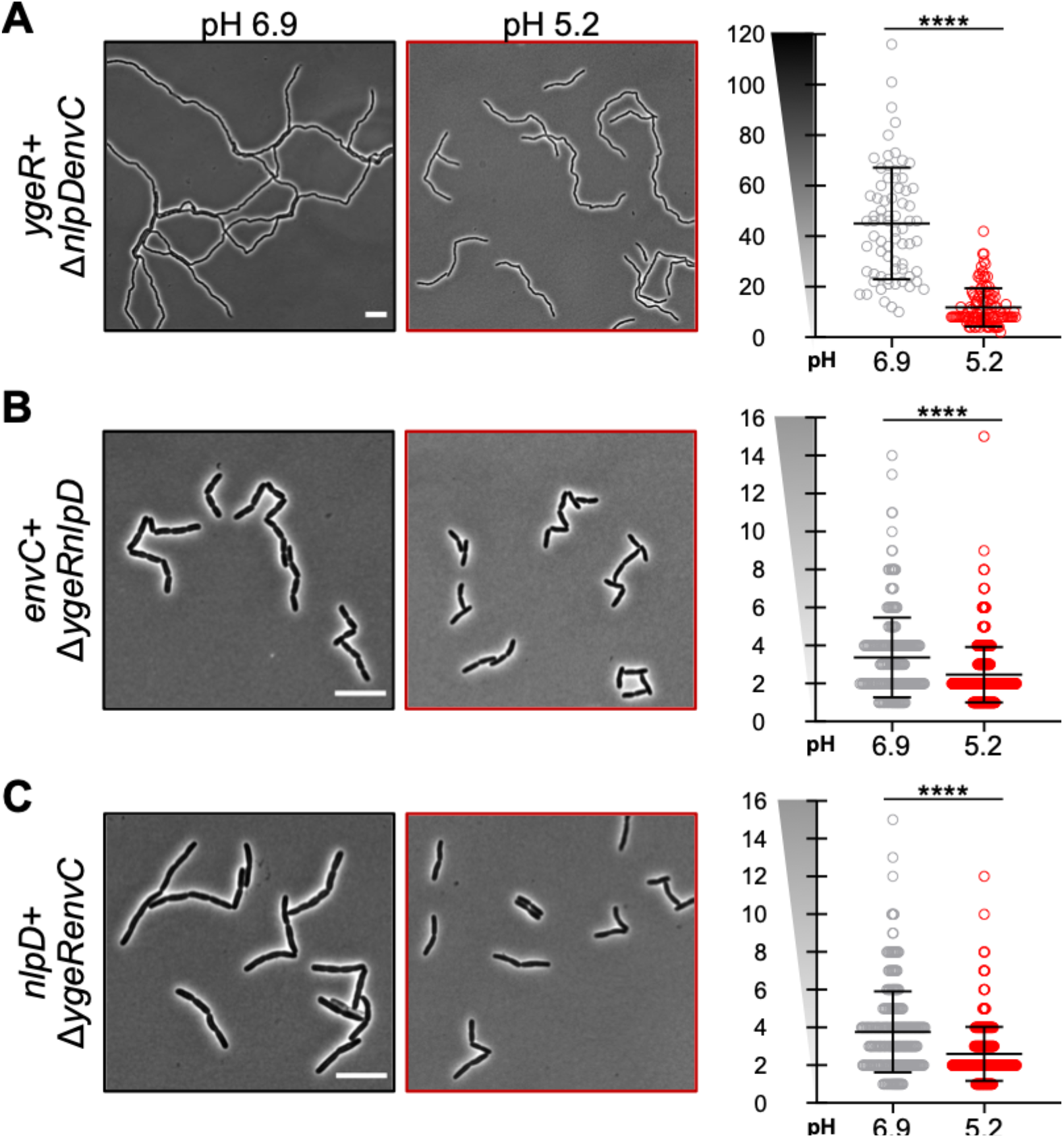
LtyM-domain protein activation at acidic pH. Representative micrographs (left) and quantification of cell chaining (right) for Δ*nlpDenvC* (EAM1372, A), Δ*ygeRnlpD* (EAM1516, B), Δ*ygeRenvC* (EAM1540, C) during steady state growth in LB medium buffered to pH 6.9 or 5.2. Cells were grown to mid-exponential phase (OD_600_ ~ 0.2-0.6) at the indicated pH, sub-cultured into the same medium at an OD_600_ ~ 0.005, then sampled at OD_600_ ~ 0.1-0.2 for live cell microscopy. Points indicate individual chain length measurements from at least two independent experiments, and error bars denote standard deviation. Y-axis gradient emphasizes the difference in scale used between mutants. Significance was determined by Kolmogorov-Smirnov test with a threshold of statistical significance set to p < 0.01. Scale bar denotes 10 μm.

### The LytM-domain protein YgeR contributes to amidase-dependent cell separation in acidic medium

Several models could explain the Δ*nlpDenvC* mutant’s enhanced cell separation in acidic medium. First, lysis of cells internal to the chain may be sufficient to separate cells and reduce overall chain length at low pH. However, unlike their counterparts at pH 6.9, few Δ*nlpDenvC* chains terminated with lysed cells at pH 5.2 (**Fig. 6B**), as would be expected if chain breakage was the predominant mechanism for cell separation (44). Alternatively, acidic pH may stimulate AmiB or AmiC either directly or through an interaction with an unknown activator.

Apart from EnvC and NlpD, *E. coli* produces two additional proteins with LtyM domains: 1) YgeR, a lipoprotein of unknown function, and 2) MepM, a DD-endopeptidase also known as YebA (**Fig. 6A**). Inactivation of either or both genes only modestly enhances the chaining of Δ*nlpDenvC* mutants during growth in standard culture conditions (37), suggesting a minor role in cell separation in these conditions. To investigate whether either YgeR or MepM play a more predominant role in cell separation in acidic medium, we inactivated each gene in the Δ*nlpDenvC* background and compared chaining across pH conditions.

Loss of YgeR, but not MepM, abolished cell separation of Δ*nlpDenvC* cells in acidic medium. Δ*nlpDenvCygeR* cells formed long chains of >40 cells in length at both neutral and acidic pH. Δ*nlpDenvCmepM* cells, on the other hand, displayed a similar cell separation pattern as the Δ*nlpDenvC* parental strain with enhanced separation in acidic medium (**Fig. 6B**). Although its absence did not affect pH-dependent chaining, loss of MepM dramatically altered cell morphology at neutral pH. Δ*nlpDenvCmepM* chains exhibit atypical bulges and twists in neutral medium, but these morphological aberrations are largely absent when cells were cultured at pH 5.2. These data implicate MepM in the preservation of normal rod shape during growth in neutral pH medium.

### Promiscuous amidase activation by LytM-domain proteins in acidic medium

Although YgeR contributes to cell separation in acidic medium, our data demonstrates this protein is not the sole amidase activator in this condition. Δ*nlpDenvC* cells still form short chains greater in length than the *amiB+ ΔamiAC* mutant in acidic medium (**Fig. 2, 6B, 7A; SI Appendix, Table S2**). Moreover, single Δ*ygeR* mutants separate normally when cultured at low pH (**SI Appendix, Fig. S7A**). Together, this evidence suggests EnvC and/or NlpD may also have heightened activation capacity in acidic medium. To isolate the pH sensitivity of each activator, we generated additional mutants that only produce a single LytM activator and compared cell chaining across pH conditions.

Similar to the *ygeR*+ Δ*nlpDenvC* mutant, cells only expressing *envC* (Δ*nlpDygeR*) or *nlpD* (Δ*ygeRenvC*) also exhibited reductions in chaining in acidic medium. Both mutants formed short chains of 3-4 cells in length when cultured in neutral medium but existed predominately as single cells or pairs of cells at low pH (**Fig. 7; SI Appendix, Table S2**). These data indicate all three LtyM-domain proteins possess heightened amidase activation capacity in acidic growth conditions.

Our results imply heightened amidase activity in acidic medium could be a result of promiscuous activation by LtyM-domain proteins, as opposed to cognate activation between specific amidase and activator pairs as previously demonstrated *in vitro* at neutral pH (36). To differentiate between these possibilities, we inactivated each LtyM-domain protein in the *amiB*+ Δ*amiAC* background, which exhibits the most dramatic pH-dependent cell chaining phenotype (**Fig. 2**). If multiple LytM-domain proteins stimulate AmiB in acidic medium, as our model predicts, then we should still observe low pH-dependent reductions in chaining.

In accordance with promiscuous activation, inactivation of YgeR, EnvC, or NlpD alone did not abolish the pH sensitivity of *amiB*+ Δ*amiAC* cells (**Fig. 8A, B; SI Appendix, Table S2**). Strikingly, loss of EnvC—the cognate activator for AmiB at neutral pH (36)— only had a modest effect on chain length (Δ*amiAC*, mean of 4 ± 1 cells per chain; Δ*amiACenvC*, mean of 6 ± 1 cells per chain). Loss of NlpD, in contrast, had the strongest effect *(ΔamiACnlpD*, mean of 8 ± 1 cells per chain). To independently verify a dominant role for NlpD in AmiB activation at low pH, we compared cell lysis during *amiB* overexpression for mutants defective for a single LytM-domain protein. Cells defective for EnvC or YgeR exhibited similar lysis kinetics to the parental strain upon *amiB* overexpression in acidic medium. Although we note the Δ*envC* mutant has a significant growth rate defect, lysis kinetics during *amiB* overexpression remain unchanged. Loss of NlpD, in contrast, alleviated toxicity (**Fig. 8C**). Overall, our findings argue acidic conditions stimulate AmiB through multiple LtyM-domain activators with NlpD having the greatest individual effect (**Fig. 8D**).

**Fig. 8.**
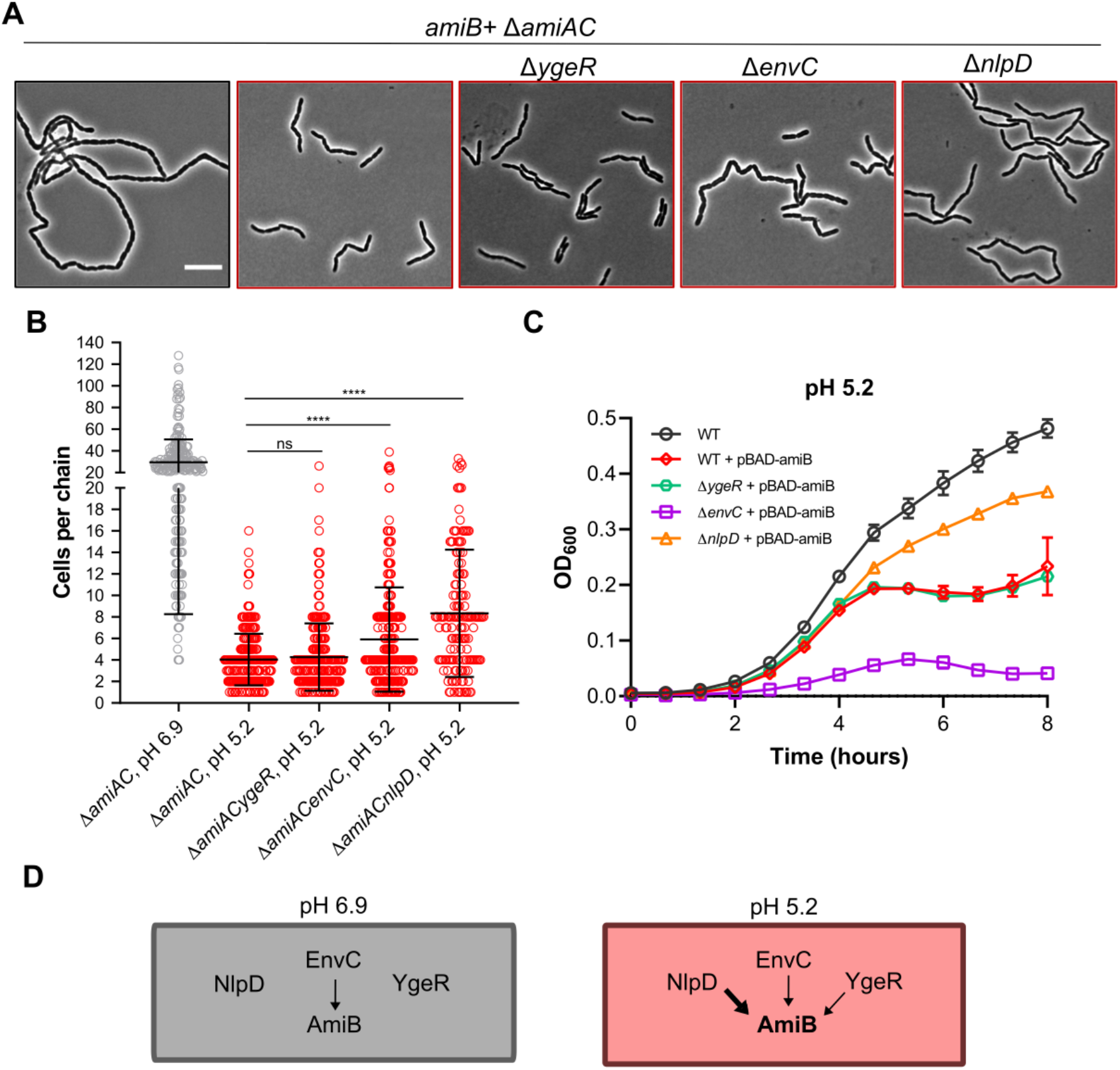
Promiscuous AmiB activation by LytM-domain proteins at acidic pH. A) Representative micrographs for Δ*amiAC* (EAM1000), Δ*amiACygeR* (EAM1549), Δ*amiACenvC* (EAM1551), and Δ*amiACnlpD* (EAM1553) during steady state growth in LB medium buffered to pH 6.9 or 5.2. Cells were grown to mid-exponential phase (OD_600_ ~ 0.2-0.6) at the indicated pH, sub-cultured into the same medium at an OD_600_ ~ 0.005, then sampled at OD_600_ ~ 0.1-0.2 for live cell microscopy. Scale bar denotes 10 μm. B) Quantification of cell chaining for mutants described in panel A. Points indicate individual chain length measurements from at least two independent experiments, and error bars denote standard deviation. Significance was determined by Kolmogorov-Smirnov test with Dunn’s correction for multiple comparison with a threshold of statistical significance set to p < 0.01. C) Growth curves of wild type (MG1655), Δ*ygeR* (EAM1514), Δ*envC* (EAM1370), and Δ*nlpD* (EAM1364) overexpressing *amiB* from a plasmid in the presence or absence of inducer (0.04% arabinose) when cultured at pH 5.2. Cells were grown to mid-exponential phase (OD_600_ ~ 0.2-0.6) in LB medium at pH 6.9, then sub-cultured into 96-well plates at an OD_600_ ~ 0.005 in LB medium buffered to 5.2. Cells were grown with aeration at 37 °C, and the optical density was measured every 10 minutes. Curves represent the average ± standard deviation of two independent experiments. D) Model of AmiB activity across pH environments. At neutral pH, EnvC specifically activates AmiB but overall AmiB activity remains low and insufficient for cell separation. At acidic pH, AmiB is activated by all three LtyM-domain proteins with NlpD playing a predominant role in stimulation.

### Computational analysis indicates pH affects amidase-activator interface

We next sought to understand the mechanistic basis for non-cognate AmiB stimulation in acidic medium. Structural modeling and genetic studies suggest the LytM activators stimulate their partner amidase(s) through a direct interaction between their degenerate (non-catalytic) LtyM domain and the target amidase’s autoinhibitory helix, resulting in the displacement of the helix from the enzyme’s active site (34). The proposed interaction interface is highly charged, implying electrostatics drive the interaction between these domains. The degenerate LtyM domains are positively charged (isoelectric point between 9.23 to 9.57 for EnvC, NlpD, and YgeR), whereas the autoinhibitory helices are negatively charged (isoelectric point between 4.28 and 5.63 for AmiA, AmiB, and AmiC). Consistent with an interaction based on electrostatics, charge inversions in the autoinhibitory helix of AmiB or the LtyM domain of EnvC impact amidase activity *in vivo* and *in vitro* (33, 34). With these data in mind, we hypothesized that acidic pH may modulate the amidase-activator interaction through one or both of the following mechanisms: 1) increasing electrostatic complementation between amidase and activator at the interaction interface and 2) destabilizing the affinity of the amidase autoinhibitory helix for the enzyme’s active site.

To examine the effect of environmental pH on electrostatic complementation, we modeled the LtyM domains of EnvC, NlpD, and YgeR and the autoinhibitory helices of AmiA, AmiB, and AmiB at pH 6.9 and 5.2. (see **Materials and Methods**) We generated two models: one leveraged a docked structure of AmiB with EnvC that is supported by mutational analyses of both proteins (33, 34) and structural studies of the catalytic cleft of the *S. aureus* LytM protein (52) (**Fig. 9A**); the second is a structure-based holistic analysis, which is agnostic to the specific LytM domain binding cleft prediction (**SI Appendix, Fig. S8**).

**Fig. 9.**
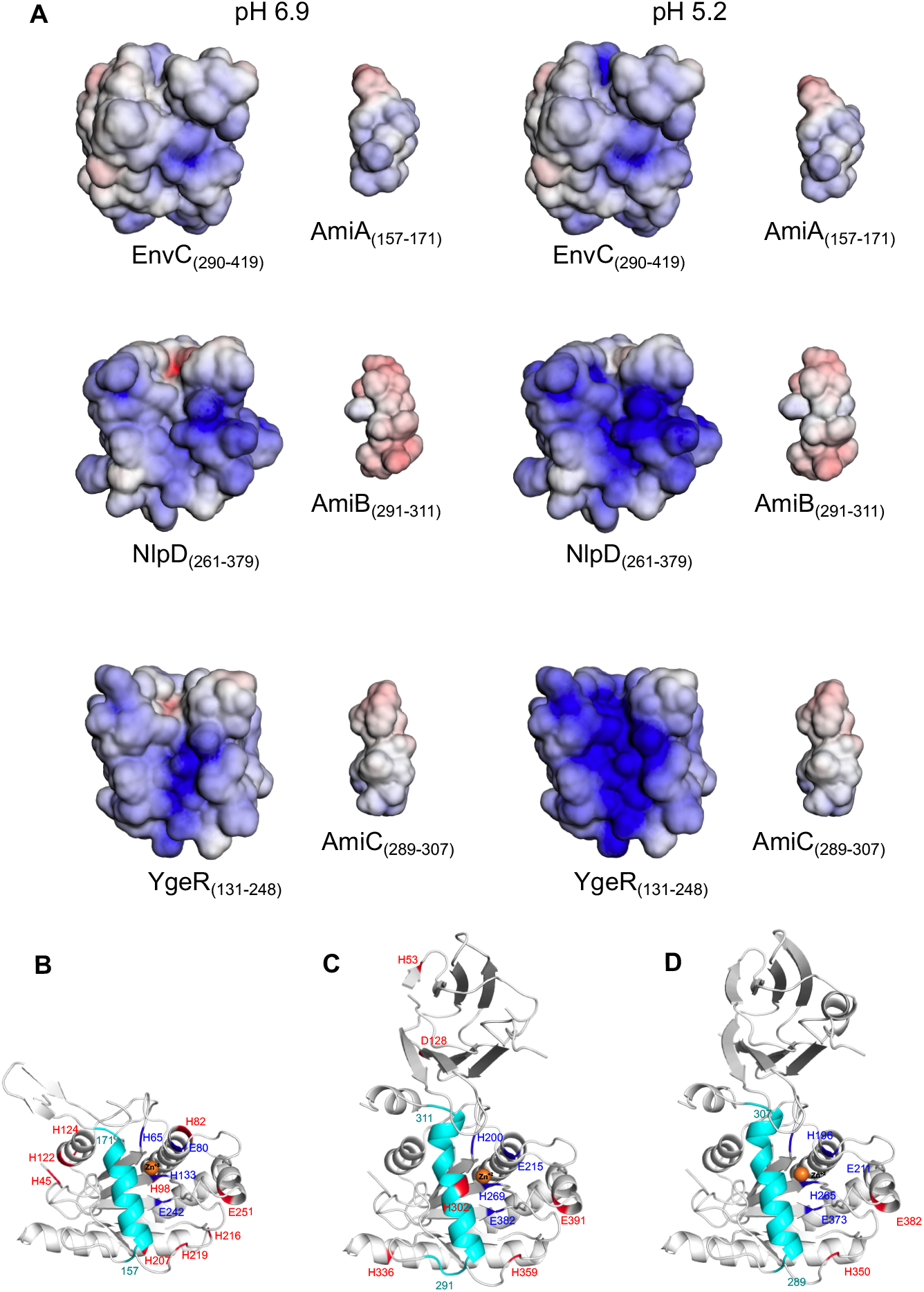
Computational analysis of electrostatic potentials and protonation state shifting residues of LytM and Ami proteins from pH 6.9 to 5.2. A) Minimum (−5 kT/e; red) and maximum (5 kT/e; blue) electrostatic potentials of the LytM domains of EnvC (residues 290-419), NlpD (residues 261-379), YgeR (residues 131-248), AmiA (157-171), AmiB (291-311) and AmiC (289-307), calculated using the APBS-PDB2PQR software (73, 74) at pH 6.9 and 5.2. B-D) Residues that change protonation state from pH 6.9 to 5.2 are shown and highlighted in red for AmiA (residues 41-276) (B), AmiB (residues 19-416) (C) and AmiC (residues 30-408) (D) structures. Zn^+2^ is shown as orange sphere, active site in dark blue and LytM domain-interacting helices in cyan (AmiA: residues 157-171; AmiB: residues 291-311; AmiC: residues 289-307). The figure was generated with PyMOL (78).

The results of the computational models reinforce our genetic data that AmiB can interact with all three activators at acidic pH. AmiB maintains a high negative charge across its autoinhibitory helix at both pH 6.9 and 5.2. On the other hand, all three of the amidase activators’ LytM domains become more positively charged at pH 5.2. The surface-exposed residues in the YgeR LytM cleft displays the greatest positive shift in electrostatic potential (**Fig. 9A**). It is tempting to speculate that the reduced surface charge at neutral pH does not favor the interaction of between YgeR and AmiB, thus explaining why YgeR does not significantly contribute to cell separation in this condition (**Fig. 6**). Of the three activators, the surface charges of NlpD and YgeR’s LytM clefts were most complementary to the AmiB autoinhibitory helix (**Fig. 9A**).

We also modeled the protonation state of residues in AmiA, AmiB, and AmiC at pH 6.9 and 5.2 (see Materials and Methods). Strikingly, this analysis revealed a putative pH-sensitive histidine (H302) in AmiB’s autoinhibitory helix that abuts the enzyme’s catalytic core (Fig. 9C) and has previously been implicated in AmiB autoinhibition (33). AmiA and AmiC, in contrast, are not predicted to possess pH-sensitive residues within their autoinhibitory helices (Fig. 9B, D). All-in-all, our computational analysis indicates that low pH-dependent AmiB activation may be driven by a combination of favorable electrostatic complementation with non-cognate LtyM activator proteins and the destabilization of its own autoinhibitory helix.

## DISCUSSION

Here, we report two of *E. coli*’s three LytC-type-*N*-acetylmuramoyl-*L*-alanine amidases exhibit pH-dependent changes in activity. While AmiB or AmiC alone cannot support normal cell separation or cell envelope integrity in neutral medium, low pH increases their activity by at least 2-fold *in vivo* (**Fig. 3, 5**), permitting formation of short chains of two to four cells and single bacteria (**Fig. 2**). The pH-dependence of amidase activity is most stark for AmiB. Our genetic and cytological data indicate this enzyme has only minimal activity *in vivo* in neutral medium when it is specifically stimulated by its cognate LytM-domain protein activator EnvC (36). In acidic medium, promiscuous activation by three LtyM-domain proteins—EnvC, NlpD, and YgeR—augments AmiB’s ability to support cell separation (**Fig. 8D**). This finding builds upon our previous work (21) and challenges the perception of redundancy within periplasmic PG enzymes. Instead of being redundant, PG enzymes exhibit overlapping activities with some specialized for distinct environmental conditions. This mechanism may ensure robust cell wall construction across environmental niches in which cells naturally grow outside the laboratory (26).

### LtyM-domain proteins enhance amidase activity in acidic environments

Our *in vitro* and *in vivo* data support a model in which acidic conditions stimulate AmiB and AmiC through the LtyM-domain proteins EnvC, NlpD, and YgeR. Importantly, our work firmly establishes YgeR as a condition-dependent amidase activator. While EnvC and NlpD have long been implicated in amidase activation in *E. coli* (32, 34–38, 50, 51, 53, 54) and other Enterobacteriaceae (55, 56), it remained unclear whether YgeR or MepM stimulate amidase activity like their counterparts. Loss of either factor only modestly exacerbates the cell separation phenotypes of mutants defective for EnvC and/or NlpD at neutral pH (37). Here, we demonstrate YgeR—but not MepM— preferentially participates in cell separation in acidic conditions (**Fig. 6**), possibly due to a more favorable electrostatic complementation for amidase activation in this environment (**Fig. 9A**). Similar to the *bone fide* amidase regulators EnvC and NlpD (34), YgeR does not likely possess any intrinsic hydrolytic activity. The LtyM domain of YgeR contains substitutions at key amino acid residues implicated in zinc coordination and hydrolytic activity among LytM family metallopeptidases (e.g., LytM of *Staphylococcus aureus*) (**SI Appendix, Fig. S7B**) (57–61). Nevertheless, future studies will be required to confirm both the absence of YgeR hydrolase activity and its ability to promote amidase activity *in vitro*. MepM, in contrast, shares conserved catalytic residues with hydrolytic members of the LtyM family and accordingly possesses metalloendopeptidase activity (7). Curiously, although MepM does not participate in cell separation, our data indicate that its activity may still be pH sensitive, at least in the absence of EnvC and NlpD. Inactivation of MepM confers morphological aberrations in neutral, but not acidic, medium (**Fig. 6**). Future work will be necessary to uncover the basis for this condition-dependent phenotype.

Our findings also challenge the prevailing model of amidase activation in which the activity of an individual amidase is specifically stimulated by a cognate LytM-domain regulator *in vivo* (33, 34, 36–38). Although collectively required for low pH-mediated cell separation (**Fig. 6**), neither EnvC, NlpD, or YgeR alone account for full AmiB activation during growth in acidic medium (**Fig. 8A, B**). Unexpectedly, NlpD has the greatest individual impact on AmiB hyperactivation at low pH: loss of NlpD increases the average chain length of *amiB*+ Δ*amiAC* and relieves lysis during AmiB overproduction in acidic medium (**Fig. 8A-C**). Consistent with NlpD serving as the predominant AmiB activator, our computational analysis predicts NlpD to have the most favorable electrostatic potential for AmiB interaction (**Fig. 9A**). Loss of EnvC, in contrast, only minimally affects these phenotypes despite its critical role in AmiB activation at neutral pH (36). We note that there is precedent for non-specific amidase activation by LtyM-domain proteins in the Enterobacteriaceae. In *Vibrio cholerae*, both EnvC and NlpD modulate the activity of AmiB, the pathogen’s sole amidase, and NlpD is preferentially required for intestinal colonization in an infant mouse model and during exposure to bile salts (56).

It remains unclear whether hyperactivation of EnvC, NlpD, and YgeR in acidic medium *in vivo* is solely a product of enhanced affinity for their substrates (**Fig. 9**) or requires additional upstream signals. Work from other groups suggests the ability of EnvC and NlpD to stimulate the amidases is tied to the activity of the cell division apparatus, albeit through distinct mechanisms. EnvC activation is dependent on the FtsEX complex (35, 62), an ABC transporter implicated in constriction initiation, whereas NlpD activation requires YraP, a newly discovered division protein of unknown function (38). Although the YgeR activation cascade remains unknown, this protein is also modestly enriched at midcell (37) in proximity to the division machinery. Consistent with a role for the cell division apparatus in the stimulation of amidase activity, we recently demonstrated that acidic pH promotes activation of cytokinesis in *E. coli*. Specifically, the cell division protein “trigger” FtsN hyperaccumulates at midcell during growth at low pH and stimulates cytokinesis at a reduced cell volume in this condition (63). It is tempting to speculate hyperactivation of division at low pH increases the activity of the LytM-domain proteins. AmiB and AmiC midcell localization is likewise responsive to cell division activation and in particular to FtsN accumulation at the cytokinetic ring. While both EnvC and NlpD accumulate at midcell prior to constriction initiation (32, 62), AmiB and AmiC require active septal cell wall synthesis and midcell localization of the terminal cell division protein FtsN for their own recruitment to the cytokinetic machinery (14, 32). It is conceivable low pH-mediated septal enrichment of FtsN leads to hyperaccumulation of AmiB and AmiC at the cytokinetic ring, increasing the proportion of enzymes available for activation by the LtyM-domain proteins. FtsN-mediated amidase recruitment may be self-reinforcing: medial FtsN localization itself is enhanced by a C-terminal periplasmic SPOR domain, which binds denuded glycans—the products of amidase activity (47, 48). Compellingly, this model may also explain why AmiA appears insensitive to pH. Unlike AmiB and AmiC, AmiA does not exhibit midcell localization and is instead peripherally distributed (14). Unfortunately, two obstacles made it difficult to conclusively compare AmiB and AmiC localization across pH conditions: 1) pH-dependent changes in intensity of periplasmic fluorescent proteins fusions, and 2) loss of amidase septal localization during the fixation process in our hands. Support for this model thus awaits further investigation.

### Widespread condition-dependent activity of extracytoplasmic cell wall enzymes

Our present work adds *E. coli* AmiB, AmiC, and the three LtyM-domain amidase activators to a growing list of pH-sensitive cell wall enzymes and regulators (19–21, 24, 63–67). Remarkably, while *E. coli* PG composition remains nearly unchanged during growth in different pH environments (19), pH “specialist” enzymes are present in every major category of its cell wall autolysins (19, 21, 68). From a technical standpoint, widespread conditional activity of PG enzymes underscores the need to report and standardize culture conditions when assessing the growth and morphological phenotypes of cell wall mutants. Differences in culture conditions and growth phase of sampling may explain discrepancies between chaining phenotypes for amidase mutants in previous studies (5, 40). More broadly, conditional activation of PG enzymes supports the hypothesis that the active repertoire of extracytoplasmic PG enzymes in a given cell changes based on the characteristics of its growth environment (9, 26). Attractively, this model explains how the extracytoplasmic reactions in PG synthesis remain robust in spite of the unstable chemical and physical properties of the periplasm (28). It is tempting to speculate that specialization of PG enzymes evolved to allow environmental generalists like *E. coli* to grow across a wide range of conditions.

## MATERIALS AND METHODS

### Bacterial strains, plasmids, and growth conditions

Unless otherwise indicated, all chemicals, media components, and antibiotics are purchased from Sigma Aldrich (St. Louis, MO). All deletion alleles, with the exception of the Δ*amiB::kan* (a gift from Daniel Kahne (40)) and Δ*ygeR::cm* (generated by lambda red recombineering (39, 69)), were originally supplied by the Coli Genetic Stock from the Keio collection (39) and transduced via P1 transduction into the MG1655 background, referred to as ‘wild type’ (WT) in the text. Transductants have been confirmed with diagnostic PCR and sequencing. Unless otherwise indicated, strains are grown in lysogeny broth (LB) medium (1% tryptone, 1% NaCl, 0.5% yeast extract) supplemented with 100 mM filter-sterilized MMT buffer (1:2:2 molar ratio of D,L-malic acid, MES, and Tris base) to vary media pH values. When selection was necessary, cultures were supplemented with 50 μg/ml kanamycin (Kan), 25-100 μg/ml ampicillin (Amp), or 15-30 μg/ml chloramphenicol. Cells were grown at 37°C in glass culture tubes shaking at 200 rpm for aeration unless otherwise stated in the figure legends.

### Growth rate measurements

Strains were grown from single colonies in glass culture tubes in LB + MMT buffer (pH 6.9) to mid-log phase (OD_600_ ~0.2-0.6), pelleted, and re-suspended to an OD_600_ 1.0 (~1 x 10^9^ CFU/mL). With the exception of Δ*amiABC*, cells were diluted and inoculated into fresh LB + MMT buffer at various pH values in 12-well (1 mL final volume) or 96-well plates (150 μl final volume) at 1 x 10^3^ CFU/mL. Plates were grown at 37°C shaking for 20 hours in a BioTek Eon microtiter plate reader, measuring the OD_600_ of each well every ten minutes. For the Δ*amiABC* mutant (which clumps extensively and cannot be reliably measured in a plate reader), cells were sub-cultured into glass test tubes (7 mL final volume), sampled at various time points, and vortexed for 30 seconds prior to measuring OD_600_. Doublings per hour was calculated by least squares fitting of early exponential growth (OD_600_ between 0.005-0.1). At least two biological replicates were performed.

### Microscopy

Phase contrast and fluorescence images of live or fixed cells were acquired from samples on 1% agarose/phosphate buffered saline or H_2_O pads with a Nikon TiE inverted microscope equipped with a 100X Plan N (N.A. = 1.25) objective (Nikon), SOLA SE Light Engine (Lumencor), heated control chamber (OKO Labs), and ORCA-Flash4.0 sCMOS camera (Hammamatsu Photonics). Filter sets for fluorescence were purchased from Chroma Technology Corporation. Nikon Elements software (Nikon Instruments) was used for image capture.

### Cell chaining measurements

Cells were initially cultured from a single colony in LB medium + MMT buffer (pH 5.2 or 6.9) to the exponential growth phase (OD_600_ ~0.2-0.6). Cultures were then back-diluted into fresh media to an OD_600_ = 0.005 and grown to early exponential phase (OD_600_ ~0.1-0.2). When necessary, 0.5 μM Mitotracker (Thermo Fisher, Waltham, MA) was added to the cultures upon back dilution in order to visualize septa in chains without clear constriction sites. Live cells were washed two times in 1x phosphate buffered saline then mounted onto 1% agarose (in either H_2_O or phosphate buffered saline) pads and allowed to dry. Cells (500 μL) were fixed by adding 20 μL of 1M NaPO4 (pH 7.4) and 100 μL of fixative (16% paraformaldehyde + 8% glutaraldehyde). Samples were incubated at room temperature for 15 minutes, then on ice for 30 minutes. Fixed cells were pelleted, washed three times in 1 mL 1X phosphate buffered saline (PBS, pH 7.4) then resuspended in GTE buffer (glucose-tris-EDTA) and stored at 4°C. Images were acquired for analysis within 48 hours of fixation. Chain length (i.e., number of cells per chain) was determined by eye or by detection of septa in MicrobeJ (70). We only counted chains that were within the micrograph field of view, so the presented data underestimates chain length. At least 50 chains across at least 2 biological replicates were counted.

### Triton-X sensitivity

For determination of minimum inhibitory concentrations, cells were grown from a single colony in LB medium + MMT buffer (pH 5.2, 6.9) at the indicated pH to mid-exponential phase (OD_600_ ~ 0.2-0.6) at 37°C with aeration and then inoculated at 1 x 10^5^ CFU/mL into the same medium in sterile 96-well plates with a range of two-fold dilutions of Triton-X (final volume, 150 μL). Plates were incubated at 37°C shaking for 20 hours before determination of the well with the lowest concentration of the antibiotic that had prevented growth by visual inspection. Three biological replicates were performed.

### Lysis curves

For overexpression experiments, wild-type cells harboring the indicated plasmids were grown from single colonies in glass culture tubes in LB + MMT buffer (pH 6.9) to mid-log phase (OD_600_ ~0.2-0.6) in the absence of inducer, pelleted, and re-suspended to an OD_600_ 1.0. Cells were diluted and inoculated into fresh LB + MMT buffer (pH 6.9 or pH 5.2) with various concentrations of inducer in 96-well plates (150 μl final volume) and glass culture tubes (5 mL final volume) at OD_600_ = 0.005. Plates were grown at 37°C shaking for 20 hours in a BioTek Eon microtiter plate reader. For morphological analysis, cells were sampled from glass culture tubes at the indicated time points, spotted on to 1% agarose pads (phosphate buffered saline), and imaged. Three biological replicates were performed.

### SPOR domain binding assay

His_6_-GFP-DamX^SPOR^ purification was performed as previously described (47) with minor modifications. Briefly, 1L of culture (BL21/pDSW1171) was grown in LB medium at 37 °C to OD_600_ ~ 0.5. Cells were transferred to 18 °C and induced by the addition of isopropyl-β-D-thiogalactopyranoside to 1 mM overnight. Cells were harvested and broken by sonication, and the His_6_-tagged protein was purified by cobalt-affinity chromatography. Purity was assessed by SDS-PAGE, and protein concentration was determined by Bradford assay using bovine serum albumin (BSA) standards.

To label the cells, 100 ng/mL of His_6_-GFP-DamX^SPOR^ and 100 ng/mL of BSA were added to ~2.5 x 10^6^ fixed cells (see “Cell chaining measurements”) in phosphate buffered saline (pH 7.4). Reactions were incubated on ice for 30 minutes, washed two times in phosphate buffered saline, and resuspended in GTE buffer. Cells were spotted on to 1% agarose pads (H_2_O) and imaged by fluorescence microscopy. Two independent biological replicates were performed. Cell segmentation and fluorescence profiles were generated in MicrobeJ (70). For each replicate, relative fluorescence intensity was normalized to the maximum value for wild-type cells at pH 6.9.

### Protonation state shifting residues of Ami proteins for pH 6.9 to 5.2 transition

The Phyre web-server (71 was used to model AmiA (residues 41-276, template PDB id: 4BIN (53)), AmiB (residues 19-416, template PDB id: 4BIN (53)) and AmiC X-ray crystal structure was downloaded from the RCSB PDB database (72) (residues 30-408, PDB id: 4BIN (53)). The PDB2PQR software (73, 74) was used for atomic charge assignments at pH 6.9 and 5.2 with the PARSE force field (75) and the PROPKA tool (76). Then atomic charges were converted to residue charges by summing the atomic charges of each residue. Residue charge changes upon pH shift were calculated by subtraction of residue charges at pH 6.9 and 5.2.

### Molecular modeling of the LytM domains of EnvC, NlpD, YgeR and MepM

The EnvC X-ray structure was downloaded from RCSB PDB database (72) (residues 278-419 PDB id: 4BH5 (34)) and the EnvC LytM domain (residues 290-419) was extracted. The 3D structures of NlpD, YgeR and MepM were modeled using the Phyre web-server (71) using PDB id: 6U2A (77) as template for all three structures. The LytM domains of NlpD (residues 261-379), YgeR (residues 131-248) and MepM (residues 287-412) were then extracted using the PyMOL software (78).

### Electrostatic potential calculations of LytM domains and Ami helices at pH 6.9 and 5.2

The LytM domain structures of EnvC, NlpD, YgeR, and MepM and their interacting helixes of the amidases (AmiA: residues 157-171; AmiB: residues 291-311, and AmiC: residues 289-307) were uploaded to the APBS-PDB2PQR web-based software (73, 74) using the PARSE force field (75)and the PROPKA tool (76) for pH 5.2 and 6.9 with N- and C-termini neutralization options. Individual electrostatic potential snapshots were recorded using the web-embedded molecular visualization tool for minimum and maximum potential limits of −5 kT/e and 5 kT/e.

## ACKNOWLEDGEMENTS

We thank David Weiss for the gift of pDSW1171, Daniel Kahne for the gift of HSC078, Corey Westfall for helpful discussions, and Navaneethan Palanisamy for technical assistance. This work was supported by NIH GM127331 to P.A.L., a Washington University Biology Summer Undergraduate Research Fellowship (BioSURF) to A. G. I., NSF graduate research fellowship DGE-1745038 to E.A.M., and a Center for Science and Engineering of Living Systems (CSELS) graduate scholar fellowship to E.A.M.

---

**Fig. S1. Effect of pH on amidase mutant growth and morphology.** A) Representative micrographs of single amidase mutants [Δ*amiA* (EAM759), Δ*amiB* (EAM1377), and Δ*amiC* (EAM763)] during steady state growth in LB medium buffered to pH 6.9 or 5.2 compared to the parental strain (MG1655). Cells were grown to midexponential phase (OD_600_ ~ 0.2-0.6) at the indicated pH, sub-cultured into the same medium at an OD_600_ ~ 0.005, then sampled and fixed at OD_600_ ~ 0.1-0.2 for microscopy. Scale bar denotes 10 μm. Quantification is shown in **Table S2**. B-D) Growth curves of single (B), double (C), and triple (D) mutants when cultured at pH 6.9 or 5.2. Cells were grown to mid-exponential phase (OD_600_ ~ 0.2-0.6) in LB medium at pH 6.9, then sub-cultured into LB medium buffered to either pH 6.9 or 5.2. Cells were grown with aeration at 37 °C. Curves represent the average ± standard deviation of at least two independent experiments.

**Fig. S2. Mutants defective for the twin arginine translocation protein export pathway phenotypically mimic the cell separation defect of Δ*amiAC* mutants.** Micrographs Δ*tatB* (EAM1397) and Δ*tatC* (EAM1399) mutants cultured at acidic (pH 5.2) and neutral (pH 6.9) medium. Images are representative of two independent experiments. Scale bar denotes 20 μm.

**Fig. S3. Complementation of amidase mutants defective for cell separation in neutral pH medium.** Micrographs and chain length quantifications for Δ*amiAC* (EAM927, A) and Δ*amiAB* (EAM1379, B) mutants harboring pBAD plasmids expressing individual amidase genes or a *thio* empty vector control. Cells were cultured to steady state in LB medium (pH 6.9) + 0.04% arabinose and sampled at OD_600_ ~ 0.1-0.2 for live cell imaging. Points indicate individual chain length measurements from two independent experiments, and error bars denote standard deviation. Scale bar denotes 20 μm.

**Fig. S4. Effect of amidase overproduction on cell growth rate.** A) Growth curves of wild-type (MG1655) cells overexpressing *amiB* from a plasmid in the presence 0.4% arabinose and cultured at either pH 6.9 or 5.2. Cells were grown to mid-exponential phase (OD_600_ ~ 0.2-0.6) in LB medium at pH 6.9, then sub-cultured into 96-well plates at an OD_600_ ~ 0.005 in LB medium buffered to either pH 6.9 or 5.2. Cells were grown with aeration at 37 °C, and the optical density was measured every 10 minutes. Curves represent the average ± standard deviation of three independent experiments. B) Representative micrographs of cell morphology and debris after six hours of growth with inducer. Scale bar denotes 5 μm. C-E) Growth curves of wild-type (MG1655) cells overexpressing *amiA* (D), *amiC* (C), or an empty vector control (E) from a plasmid in the presence or absence of arabinose (0.4%) at pH 6.9 or 5.2. Cells were grown to mid-exponential phase (OD_600_ ~ 0.2-0.6) in LB medium at pH 6.9, then sub-cultured into 96-well plates at an OD_600_ ~ 0.005 in LB medium buffered to either pH 6.9 or 5.2. Cells were grown with aeration at 37 °C, and the optical density was measured every 10 minutes. Curves represent the average ± standard deviation of three independent experiments.

**Fig. S5. His_6_-GFP-DamX^SPOR^ reports levels of denuded glycans at midcell.** A) Schematic depicting use of GFP-SPOR fusion protein to detect denuded glycans, the products of amidase activity. Representative micrographs (B) and quantification (C) for His_6_-GFP-DamX^SPOR^ binding to fixed wild-type, Δ6LT, and Δ*amiABC* cells cultured in LB medium at pH 6.9. Cells were cultured to steady state in LB medium (pH 6.9) and then fixed at OD_600_ ~ 0.1-0.2. Fixed cells were immediately labeled with 100 ng/mL of purified His_6_-GFP-DamX^SPOR^ protein for 30 minutes on ice, washed, and then visualized by fluorescence microscopy. Fluorescence intensity profiles were generated from at least 50 cells across two biological replicates and normalized to the maximum intensity of the wild type at pH 6.9. Error bars represent standard deviation. Scale bar denotes 5 μm. D) Cell length distribution for Δ6LT mutant grown to steady state in LB medium (pH 6.9). E) Representative micrographs and quantification for His_6_-GFP-DamX^SPOR^ binding to fixed Δ*amiBC* (EAM1381) cells. Cells were cultured to steady state in LB medium (pH 6.9 or 5.2) and then fixed at OD_600_ ~ 0.1-0.2. Fixed cells were immediately labeled with 100 ng/mL of purified His_6_-GFP-DamX^SPOR^ protein for 30 minutes on ice, washed, and then visualized by fluorescence microscopy. Scale bar denotes 5 μm. Fluorescence intensity profiles were generated from at least 50 cells across two biological replicates and normalized to the maximum intensity of the wild type at pH 6.9. Error bars represent standard deviation.

**Fig. S6. Loss of AmiB or AmiC restores chaining of Δ*nlpDenvC* at acidic pH.** Representative micrographs of Δ*nlpDenvCamiA* (EAM1422), Δ*nlpDamiBenvC* (EAM1485), and Δ*nlpDenvCamiC* (EAM1424) during steady state growth in LB medium buffered to pH 6.9 or 5.2. Cells were grown to mid-exponential phase (OD_600_ ~ 0.2-0.6) at the indicated pH, sub-cultured into the same medium at an OD_600_ ~ 0.005, then sampled and fixed at OD_600_ ~ 0.1-0.2 for microscopy. Scale bar denotes 20 μm.

**Fig. S7. Morphological and bioinformatic analysis of LtyM-domain factors.** A) Representative micrographs of Δ*envC* (EAM1370), Δ*nlpD* (EAM1364), and Δ*ygeR* (EAM1514) during steady state growth in LB medium buffered to pH 6.9 or 5.2. Cells were grown to mid-exponential phase (OD_600_ ~ 0.2-0.6) at the indicated pH, sub-cultured into the same medium at an OD_600_ ~ 0.005, then sampled and fixed at OD_600_ ~ 0.1-0.2 for microscopy. Scale bar denotes 10 μm. B) Amino acid alignment of LytM-domain (Pfam: Peptidase family 23 PF01551) family proteins from *E. coli* (MepM, EnvC, YgeR, and NlpD) with *S. aureus* LytM. Alignment begins at residue 312 for MepM. Residues at positions required for *S. aureus* LytM catalysis are highlighted in yellow, and residues required for *S. aureus* LytM zinc binding are denoted with an asterisk. Fraction denotes number of critical conserved residues with *S. aureus* LytM for MepM, EnvC, YgeR, and NlpD.

**Fig. S8. pH and ionic strength dependent charge/ (number of amino acids)] distributions of LytM domains and Ami helices** pH and ionic strength dependent [charge/ (number of amino acids)] distributions of LytM domains of EnvC-NlpD-YgeR and Ami helices (AmiA (157-171), AmiB (291-311) and AmiC (289-307)) were calculated by using Protein-sol heatmaps web-server by uploading respective protein structures.

**Table S1A. Bacterial strains used in this study.**

**Table S1B. Plasmids used in this study.**

**Table S2. Effect of pH on amidase mutants’ growth rate and chain length.**

