## Supplemental Material for "The active repertoire of *Escherichia coli* peptidoglycan amidases varies with physiochemical environment"

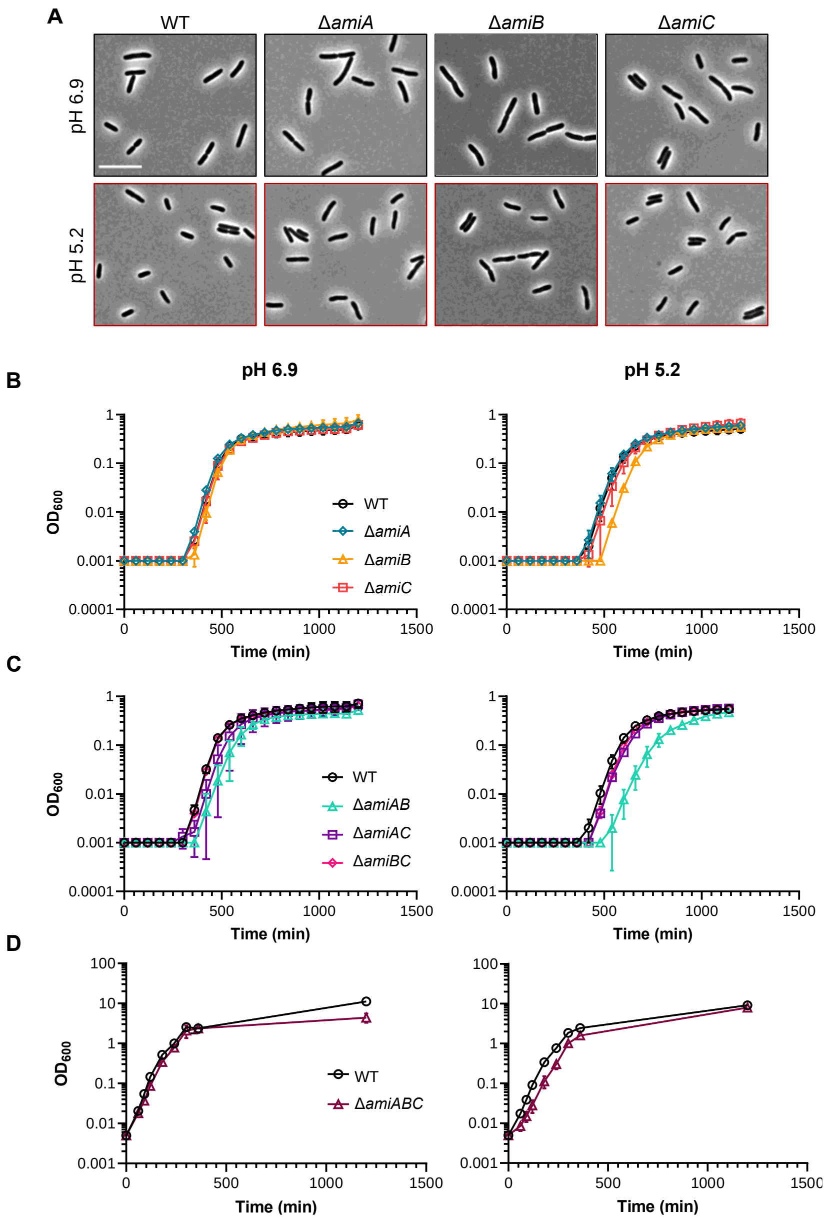


**Fig. S1. Effect of pH on amidase mutant growth and morphology.**
A) Representative micrographs of single amidase mutants [Δ*amiA* (EAM759), Δ*amiB* (EAM1377), and Δ*amiC* (EAM763)] during steady state growth in LB medium buffered to pH 6.9 or 5.2 compared to the parental strain (MG1655). Cells were grown to mid-exponential phase ($OD_{600}$ ~ 0.2-0.6) at the indicated pH, sub-cultured into the same medium at an $OD_{600}$ ~ 0.005, then sampled and fixed at $OD_{600}$ ~ 0.1-0.2 for microscopy. Scale bar denotes 10 µm. Quantification is shown in **Table S3**. B-D) Growth curves of single (B), double (C), and triple (D) mutants when cultured at pH 6.9 or 5.2. Cells were grown to mid-exponential phase ($OD_{600}$ ~ 0.2-0.6) in LB medium at pH 6.9, then sub-cultured into LB medium buffered to either pH 6.9 or 5.2. Cells were grown with aeration at 37 ºC. Curves represent the average ± standard deviation of at least two independent experiments.

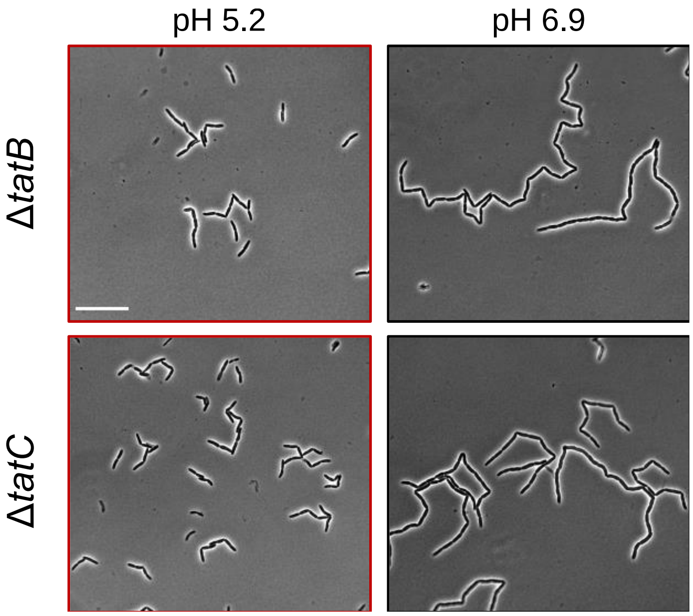


**Fig. S2. Mutants defective for the twin arginine translocation protein export pathway phenotypically mimic the cell separation defect of ∆*amiAC* mutants.** Micrographs ∆*tatB* (EAM1397) and ∆*tatC* (EAM1399) mutants cultured at acidic (pH 5.2) and neutral (pH 6.9) medium. Images are representative of two independent experiments. Scale bar denotes 20 µm.

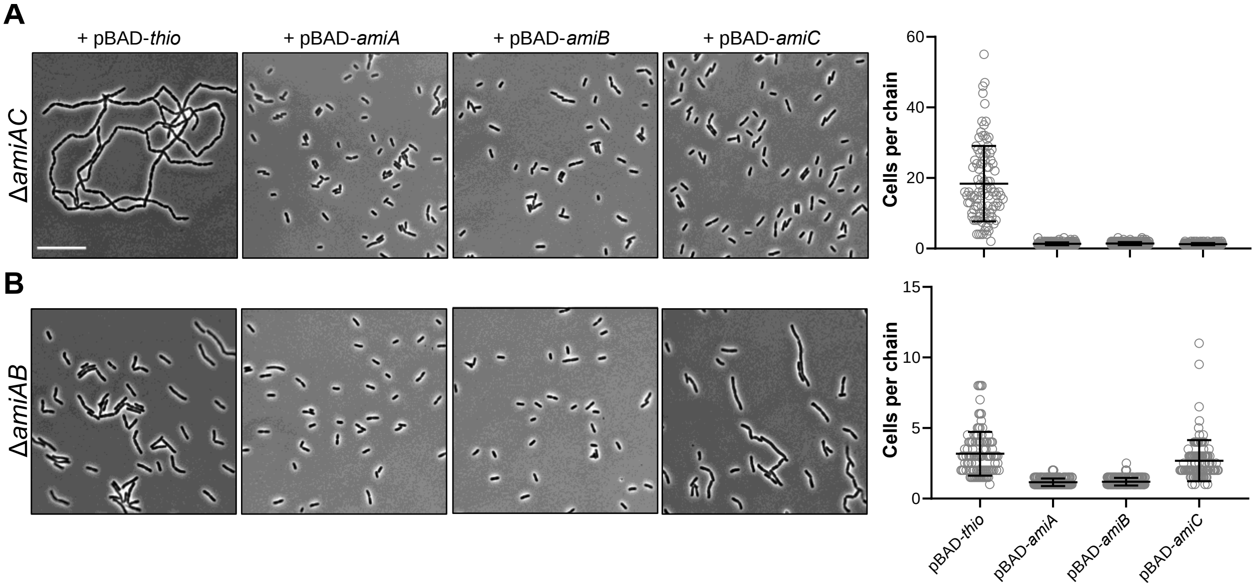


**Fig. S3. Complementation of amidase mutants defective for cell separation in neutral pH medium.**

Micrographs and chain length quantifications for Δ*amiAC* (EAM927, A) *and* Δ*amiAB* (EAM1379, B) mutants harboring pBAD plasmids expressing individual amidase genes or a *thio* empty vector control. Cells were cultured to steady state in LB medium (pH 6.9) + 0.04% arabinose and sampled at $OD_{600}$ ~ 0.1-0.2 for live cell imaging. Points indicate individual chain length measurements from two independent experiments, and error bars denote standard deviation. Scale bar denotes 20 µm.

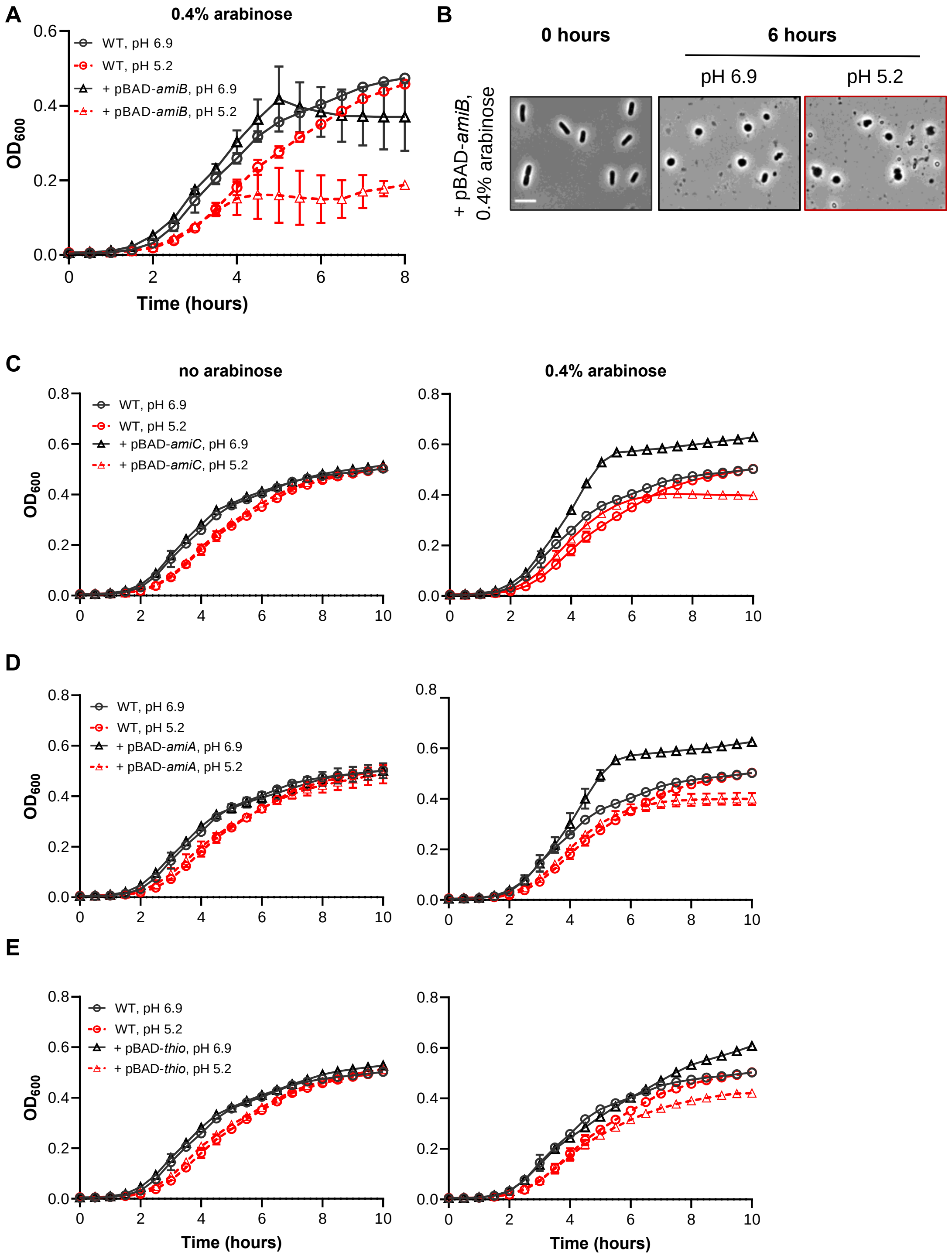


**Fig. S4. Effect of amidase overproduction on cell growth rate.**
A) Growth curves of wild-type (MG1655) cells overexpressing *amiB* from a plasmid in the presence 0.4% arabinose and cultured at either pH 6.9 or 5.2. Cells were grown to mid-exponential phase ($OD_{600}$ ~ 0.2-0.6) in LB medium at pH 6.9, then sub-cultured into 96-well plates at an $OD_{600}$ ~ 0.005 in LB medium buffered to either pH 6.9 or 5.2. Cells were grown with aeration at 37 ºC, and the optical density was measured every 10

minutes. Curves represent the average ± standard deviation of three independent experiments. B) Representative micrographs of cell morphology and debris after six hours of growth with inducer. Scale bar denotes 5 µm. C-E) Growth curves of wild-type (MG1655) cells overexpressing *amiA* (D), *amiC* (C), or an empty vector control (F) from a plasmid in the presence or absence of arabinose (0.4%) at pH 6.9 or 5.2. Cells were grown to mid-exponential phase ($OD_{600}$ ~ 0.2-0.6) in LB medium at pH 6.9, then sub-cultured into 96-well plates at an $OD_{600}$ ~ 0.005 in LB medium buffered to either pH 6.9 or 5.2. Cells were grown with aeration at 37 ºC, and the optical density was measured every 10 minutes. Curves represent the average ± standard deviation of three independent experiments.

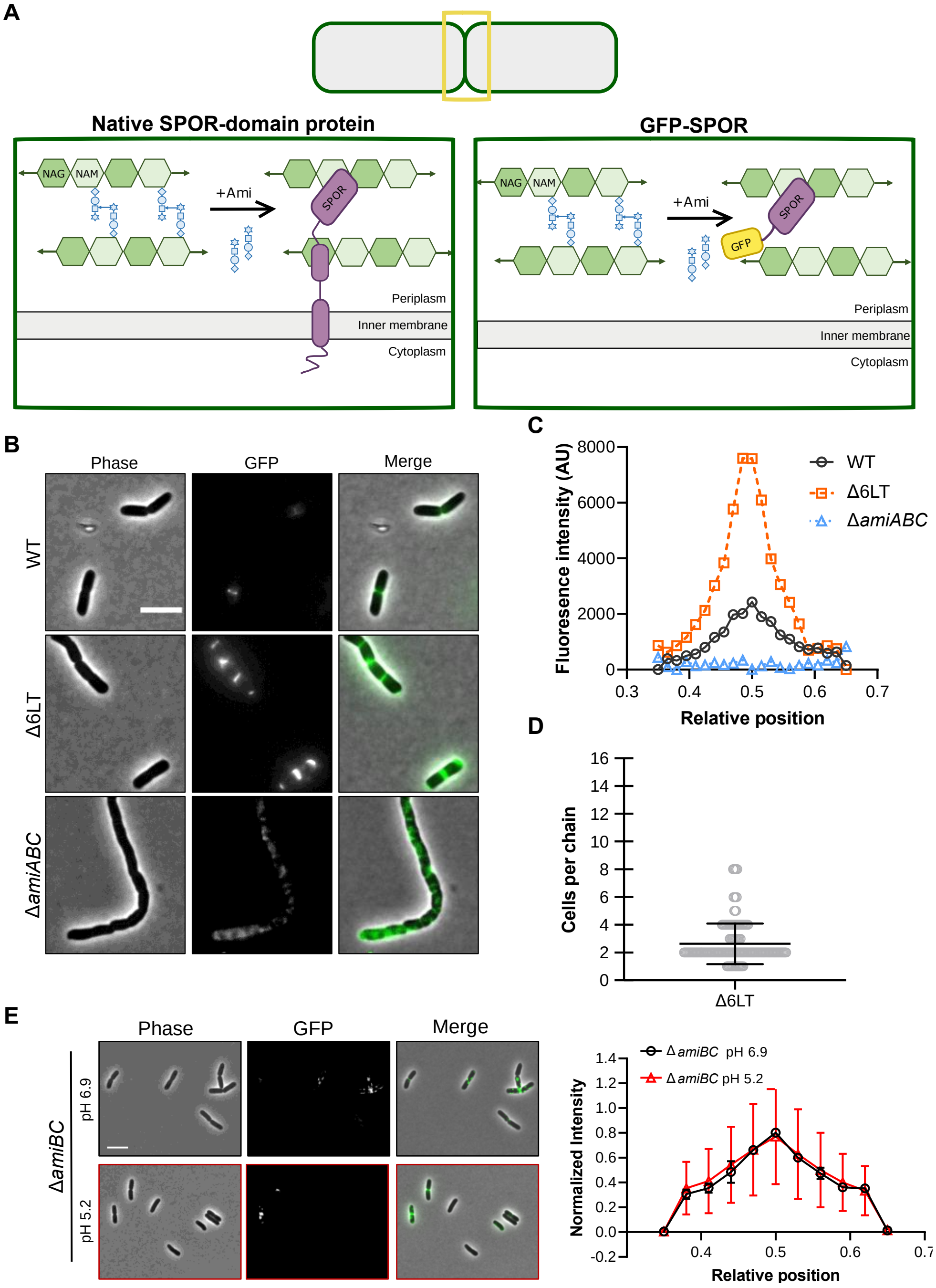


**Fig. S5. His$_6$-GFP-DamX$^{SPOR}$ reports levels of denuded glycans at midcell.**
A) Schematic depicting use of GFP-SPOR fusion protein to detect denuded glycans, the products of amidase activity. Representative micrographs (B) and quantification (C) for His$_6$-GFP-DamX$^{SPOR}$ binding to fixed wild type, Δ6LT, and Δ*amiABC* cells cultured in LB medium at pH 6.9. Cells were cultured to steady state in LB medium (pH 6.9) and then fixed at $OD_{600}$ ~ 0.1-0.2. Fixed cells were immediately labeled with 100 ng/mL of purified His$_6$-GFP-DamX$^{SPOR}$ protein for 30 minutes on ice, washed, and then visualized by fluorescence microscopy. Fluorescence intensity profiles were generated from at least 50 cells across two biological replicates and normalized to the wild type maximum intensity

at pH 6.9. Error bars represent standard deviation. Scale bar denotes 5 µm. D) Cell length distribution for ∆6LT mutant grown to steady state in LB medium (pH 6.9). E) Representative micrographs and quantification for $His_6$-GFP-$DamX^{SPOR}$ binding to fixed ∆*amiBC* (EAM138) cells. Cells were cultured to steady state in LB medium (pH 6.9 or 5.2) and then fixed at $OD_{600}$ ~ 0.1-0.2. Fixed cells were immediately labeled with 100 ng/mL of purified $His_6$-GFP-$DamX^{SPOR}$ protein for 30 minutes on ice, washed, and then visualized by fluorescence microscopy. Scale bar denotes 5 µm. Fluorescence intensity profiles were generated from at least 50 cells across two biological replicates and normalized to the wild type maximum intensity at pH 6.9. Error bars represent standard deviation.

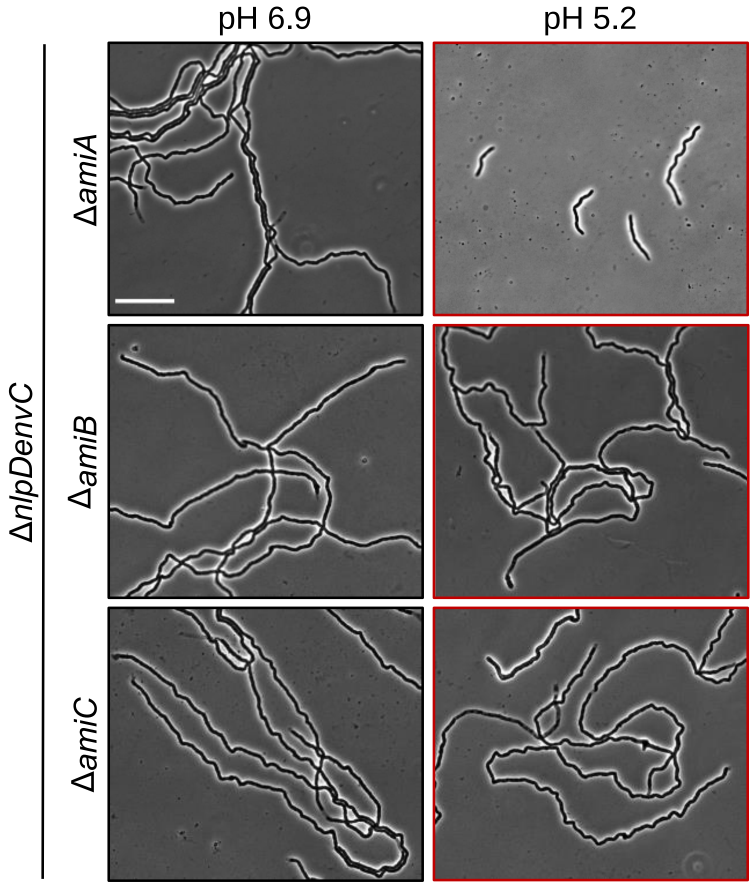


**Fig. S6. Loss of AmiB or AmiC restores chaining of Δ*nlpDenvC* at acidic pH.** Representative micrographs of Δ*nlpDenvCamiA* (EAM1422), Δ*nlpDamiBenvC* (EAM1485), and Δ*nlpDenvCamiC* (EAM1424) during steady state growth in LB medium buffered to pH 6.9 or 5.2. Cells were grown to mid-exponential phase ($OD_{600}$ ~ 0.2-0.6) at the indicated pH, sub-cultured into the same medium at an $OD_{600}$ ~ 0.005, then sampled and fixed at $OD_{600}$ ~ 0.1-0.2 for microscopy. Scale bar denotes 20 µm.

**A**

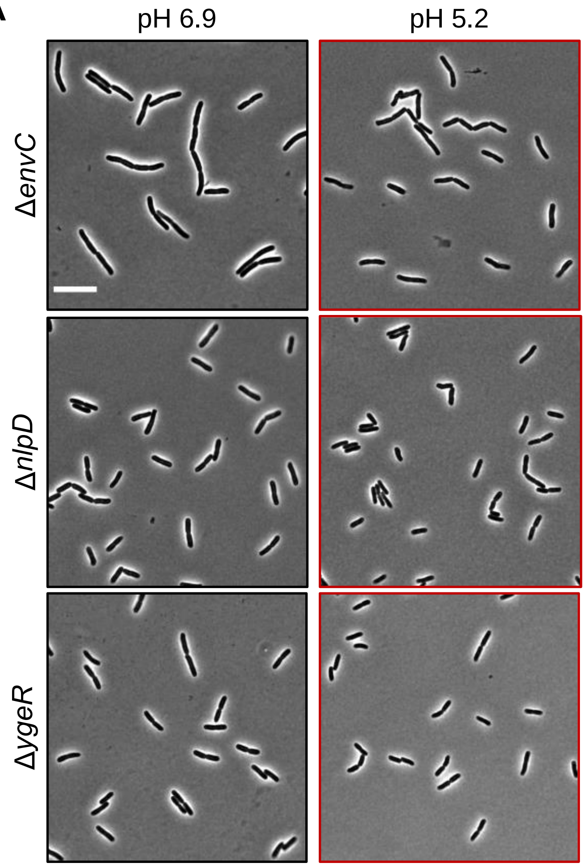


**B**

```
            *   *                                             *
MepM      APHRGVDFAMPQGTPVLSVGDGEVVVAKRSGAAG-YYVAIRHG-RSYTTRYMHLRKILVK
LtyM      GAHYGVDYAMPENSPVYSLTDGTVVQAGWSNYGGGNQVTIKEANSNNYQWYMHNNRLTVS
EnvC      -RWKGMVIGASEGTEVKAIADGRVIL-ADWLQGYGLVVVVEHG-KGDMSLYGYNQSALVS
YgeR      GGNKGIDISAPRGTPIYAAGAGKVVYVGNQLRGYGNLIMIKHS-EDYITAYAHNDTMLVN
NlpD      GGNKGIDIAGSKGQAIIATADGRVVYAGNALRGYGNLIIIKHN-DDYLSAYAHNDTMLVR

MepM      PGQKVKRGDRIALSGNTGRSTGPHLHYEVWIN----QQAVNP   (5/5)
LtyM      AGDKVKAGDQIAYSGSTGNSTAPHVHFQRMSGGIGNQYAVDP   ----
EnvC      VGSQVRAGQPIALVGSSGGQGRPSLYFEIRRQ----GQAVNP   (0/5)
YgeR      NGQSVKAGQKIATMGSTDA-ASVRLHFQIRYR----ATAIDP   (3/5)
NlpD      EQQEVKAGQKIATMGSTGT-SSTRLHFEIRYK----GKSVNP   (3/5)
```

**Fig. S7. Morphological and bioinformatic analysis of LtyM-domain factors.**
A) Representative micrographs of Δ*envC* (EAM1370), Δ*nlpD* (EAM1364), and Δ*ygeR* (EAM1514) during steady state growth in LB medium buffered to pH 6.9 or 5.2. Cells were grown to mid-exponential phase ($OD_{600}$ ~ 0.2-0.6) at the indicated pH, sub-cultured into the same medium at an $OD_{600}$ ~ 0.005, then sampled and fixed at $OD_{600}$ ~ 0.1-0.2 for microscopy. Scale bar denotes 10 µm. B) Amino acid alignment of LytM-domain (Pfam: Peptidase family 23 PF01551) family proteins from *E. coli* (MepM, EnvC, YgeR, and NlpD) with *S. aureus* LytM. Alignment begins at residue 312 for MepM. Residues at positions required for *S. aureus* LytM catalysis are highlighted in yellow, and residues required for *S. aureus* LytM zinc binding are denoted with an asterisk. Fraction denotes number of critical conserved residues with *S. aureus* LytM for MepM,

EnvC, YgeR, and NlpD. B) Amino acid alignment of LytM-domain (Pfam: Peptidase family 23 PF01551) family proteins from *E. coli* (MepM, EnvC, YgeR, and NlpD) with *S. aureus* LytM. Alignment begins at residue 312 for MepM. Residues at positions required for *S. aureus* LytM catalysis are highlighted in yellow, and residues required for *S. aureus* LytM zinc binding are denoted with an asterisk. Fraction denotes number of critical conserved residues with *S. aureus* LytM for MepM, EnvC, YgeR, and NlpD.

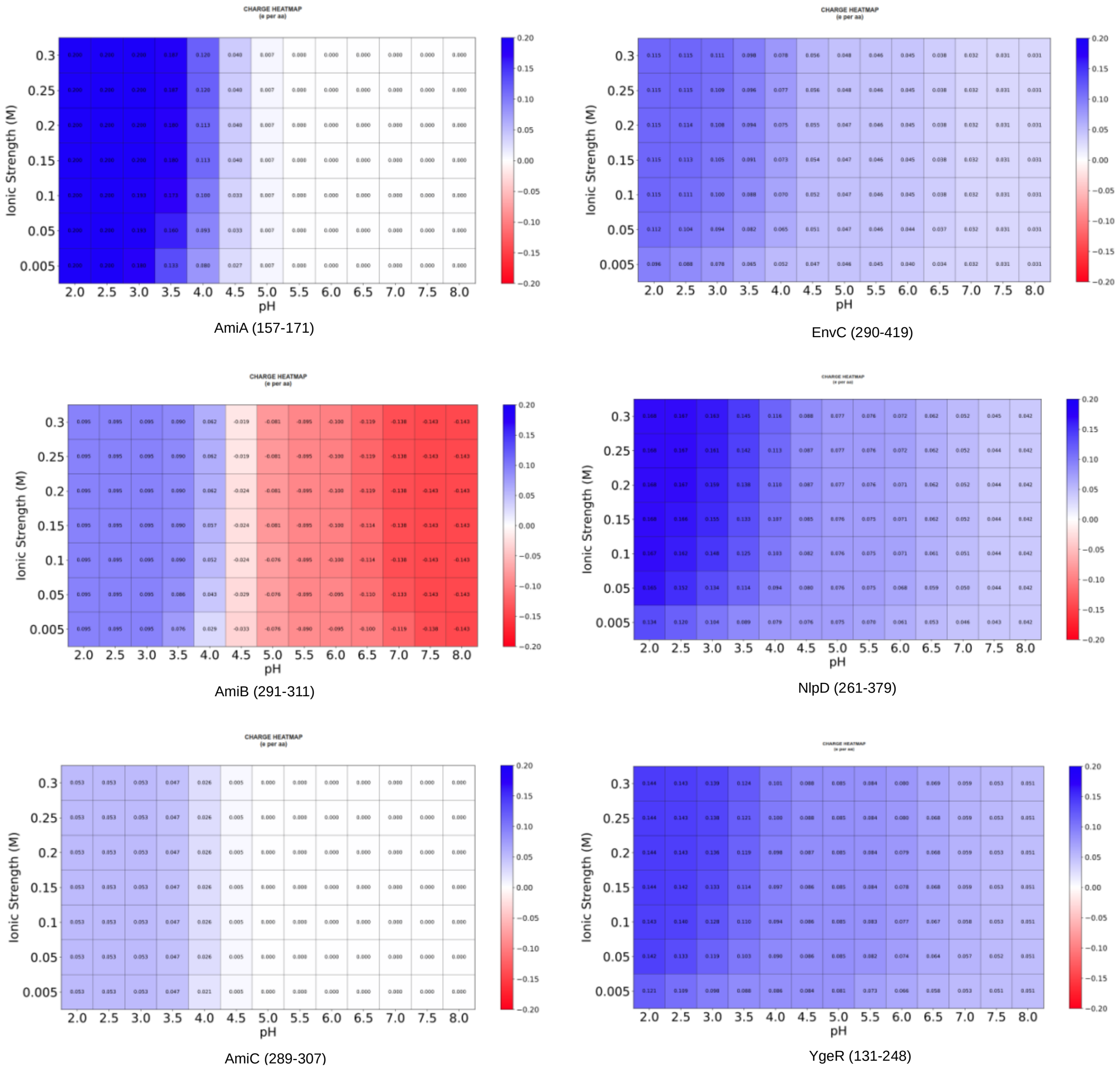


**Fig. S8. pH and ionic strength dependent charge/ (number of amino acids)] distributions of LytM domains and Ami helices.**
pH and ionic strength dependent [charge/ (number of amino acids)] distributions of LytM domains of EnvC- NlpD-YgeR and Ami helices (AmiA (157-171), AmiB (291-311) and AmiC (289-307)) were calculated by using Protein-sol heatmaps web-server by uploading respective protein structures.

**Table S1A.** Bacterial strains used in this study.

| Strain ID | Genotype | Source[a] |
|---|---|---|
| MG1655 | *rph1 ilvG rfb-50 λ- F-* | (1) |
| BL21 | *fhuA2 [lon] ompT gal [dcm] ΔhsdS* | New England Biolabs |
| HSC078 | JOE309 Δ*amiB*::*kan* | (2) |
| EAM696 | MG1655 Δ*mrcB*::*frt* | (3) |
| EAM759 | MG1655 Δ*amiA*::*kan* | (3) |
| EAM763 | MG1655 Δ*amiC*::*kan* | (3) |
| EAM806 | MG1655 Δ*nlpD*::*kan* | P1(JW2712-2[b]) x MG1655 |
| EAM924 | MG1655 Δ*amiA*::*frt* | EAM759/pCP20 |
| EAM925 | MG1655 Δ*amiC*::*frt* | EAM763/pCP20 |
| EAM927 | MG1655 Δ*amiA*::*frt* Δ*amiC*::*kan* | P1(JW4559-1[b]) x EAM924 |
| EAM1000 | MG1655 Δ*amiA*::*frt* Δ*amiC*::*frt* | EAM927/pCP20 |
| EAM1364 | MG1655 Δ*nlpD*::*frt* | EAM806/pCP20 |
| EAM1370 | MG1655 Δ*envC*::*kan* | P1(JW5646-3[b]) x MG1655 |
| EAM1372 | MG1655 Δ*nlpD*::*frt* Δ*envC*::*kan* | P1(JW5646-3[b]) x EAM1364 |
| EAM1377 | MG1655 Δ*amiB*::*kan* | P1(HSC078) x MG1655 |
| EAM1379 | MG1655 Δ*amiA*::*frt* Δ*amiB*::*kan* | P1(HSC078) x EAM924 |
| EAM1381 | MG1655 Δ*amiC*::*frt* Δ*amiB*::*kan* | P1(HSC078) x EAM763 |
| EAM1385 | MG1655 Δ*amiA*::*frt* Δ*amiC*::*frt* Δ*amiB*::*kan* | P1(HSC078) x EAM1000 |
| EAM1397 | MG1655 Δ*tatB*::*kan* | P1(JW5580-1[b]) x MG1655 |
| EAM1399 | MG1655 Δ*tatC*::*kan* | P1(JW3815-1[b]) x MG1655 |
| EAM1422 | MG1655 Δ*amiA*::*frt* Δ*envC*::*frt* Δ*nlpD*::*kan* | P1(JW2712-2[b]) x EAM1419 |
| EAM1424 | MG1655 Δ*amiC*::*frt* Δ*envC*::*frt* Δ*nlpD*::*kan* | P1(JW2712-2[b]) x EAM1418 |
| EAM1477 | MG1655 Δ*nlpD*::*frt* Δ*yebA*::*frt* Δ*envC*::*kan* | P1(JW5646-3[b]) x EAM1476 |

| | | |
|---|---|---|
| EAM1485 | MG1655 Δ*nlpD*::*frt* Δ*amiB*::*frt* Δ*envC*::*kan* | P1(JW5646-3[b]) x EAM1482 |
| EAM1514 | MG1655 Δ*ygeR*::*cm* | λ red |
| EAM1516 | MG1655 Δ*nlpD*::*frt* Δ*ygeR*::*cm* | P1(EAM1514) x EAM1364 |
| EAM1521 | MG1655 Δ*nlpD*::*frt* Δ*ygeR*::*cm* Δ*envC*::*kan* | P1(JW5646-3[b]) x EAM1516 |
| EAM1540 | MG1655 Δ*ygeR*::*cm* Δ*envC*::*kan* | P1(JW5646-3[b]) x EAM1514 |
| EAM1549 | MG1655 Δ*amiA*::*frt* Δ*amiC*::*frt* Δ*ygeR*::*cm* | P1(EAM1514) x EAM1000 |
| EAM1551 | MG1655 Δ*amiA*::*frt* Δ*amiC*::*frt* Δ*envC*::*kan* | P1(JW5646-3[b]) x EAM1000 |
| EAM1553 | MG1655 Δ*amiA*::*frt* Δ*amiC*::*frt* Δ*nlpD*::*kan* | P1(JW2712-2[b]) x EAM1000 |

[a] Strains constructed by P1 transduction are described using the shorthand: P1(donor) x recipient. Strains constructed from the removal of the Kan$^{R}$ cassette using pCP20 are indicated as follows: parental strain/pCP20 (4)
[b] Strains sourced from the Coli Genetic Stock Center (5)

**Table S1B.** Plasmids used in this study

| Plasmid ID | Genotype | Source |
|---|---|---|
| pBAD-amiA | pBAD-*amiA* | This work |
| pBAD-amiB | pBAD-*amiB* | This work |
| pBAD-amiC | pBAD-*amiC* | This work |
| pBAD-thio | pBAD-*thio* | (6) |
| pCP20 | *bla cat c*I875 *repA*(Ts) $P_R$::*flp* | (4) |
| pDSW1171 | $P_{T5}$::$His_6$-*gfp-damX*$^{SPOR}$(338–428) | (7) |

**Table S2.** Growth rates and chain lengths for amidase mutants

| Strain ID | Genotype | pH 5.2 mass doubling time (mins)[a] | pH 5.2 chain length (cells per chain)[b] | pH 6.9 mass doubling time (mins)[a] | pH 6.9 chain length (cells per chain)[b] |
|---|---|---|---|---|---|
| MG1655 | WT | 30 ± 2 | 1 [1-2] | 24 ± 1 | 1 [1-2] |
| EAM759 | Δ*amiA*::*kan* | 31 ± 1 | 1 [1-2] | 24 ± 1 | 1 [1-2] |
| EAM1377 | Δ*amiB*::*kan* | 29 ± 1 | 1 [1-2] | 24 ± 1 | 1 [1-2] |
| EAM763 | Δ*amiC*::*kan* | 31 ± 1 | 1 [1-2] | 24 ± 1 | 1 [1-2] |
| EAM1379 | Δ*amiA*::*frt* Δ*amiB*::*kan* | 42 ± 1 | 1 [1-2] | 28 ± 1 | 2 [2-4] |
| EAM927 | Δ*amiA*::*frt* Δ*amiC*::*kan* | 34 ± 1 | 4 [2-4] | 26 ± 2 | 24 [16-36] |
| EAM1381 | Δ*amiC*::*frt* Δ*amiB*::*kan* | 30 ± 1 | 1 [1-2] | 23 ± 1 | 1 [1-2] |
| EAM1385 | Δ*amiA*::*frt* Δ*amiC*::*frt* Δ*amiB*::*kan* | 34 ± 2 | 35 [20-53] | 28 ± 1 | 39 [28-58] |
| EAM1372 | Δ*nlpD*::*frt* Δ*envC*::*kan* | n.d. | 8[8-16] | n.d. | 46 [26-59] |
| EAM1521 | Δ*nlpD*::*frt* Δ*envC*::*kan* Δ*ygeR*::*cm* | n.d. | TMTC[c] | n.d. | TMTC[c] |
| EAM1516 | Δ*nlpD*::*frt* Δ*ygeR*::*cm* | n.d. | 2[2-3] | n.d. | 3[2-4] |
| EAM1540 | Δ*envC*::*kan* Δ*ygeR*::*cm* | n.d. | 2[2-3] | n.d. | 3[2-4] |
| EAM1364 | Δ*nlpD*::*frt* | n.d. | 2[1-2] | n.d. | 2[2-2] |
| EAM1370 | Δ*envC*::*kan* | n.d. | 2 [2-3] | n.d. | 3[2-4] |
| EAM1514 | Δ*ygeR*::*cm* | n.d. | 2[1-2] | n.d. | 2[2-2] |
| EAM1549 | Δ*amiA*::*frt* Δ*amiC*::*frt* Δ*ygeR*::*cm* | n.d. | 4[2-5] | n.d. | n.d. |

| | | | | | |
|---|---|---|---|---|---|
| EAM1551 | Δ*amiA*::*frt*<br>Δ*amiC*::*frt*<br>Δ*envC*::*kan* | n.d. | 4[3-8] | n.d.` | n.d |
| EAM1553 | Δ*amiA*::*frt*<br>Δ*amiC*::*frt*<br>Δ*nlpD*::*kan* | n.d. | 8[4-11] | n.d. | n.d. |

[a] mean ± SD
[b] median [25-75 percentile]
[c] TMTC denotes "to many to count"
